# Trio-binned genomes of the woodrats *Neotoma bryanti* and *N. lepida* reveal novel gene islands and rapid copy number evolution of xenobiotic metabolizing cytochrome p450 genes

**DOI:** 10.1101/2021.03.08.434435

**Authors:** Robert Greenhalgh, Matthew L. Holding, Teri J. Orr, James B. Henderson, Thomas L. Parchman, Marjorie D. Matocq, Michael D. Shapiro, M. Denise Dearing

## Abstract

The genomic architecture underlying the origins and maintenance of biodiversity is an increasingly accessible feature of species, due in large part to third-generation sequencing and novel analytical toolsets. Woodrats of the genus *Neotoma* provide a unique opportunity to study how vertebrate herbivores respond to climate change, as two sister species (*N. bryanti* and *N. lepida*) independently achieved a major dietary feat – switching to the novel and toxic food source creosote bush (*Larrea tridentata*) – in the aftermath of a natural warming event. To better understand the genetic mechanisms underlying this ability, we employed a trio binning sequencing approach with a *N. bryanti* × *N. lepida* F_1_ hybrid, resulting in phased, chromosome-level, highly complete, haploid genome assemblies for each species from one individual. Using these new assemblies, we explored the genomic architecture of three cytochrome p450 subfamilies (2A, 2B, and 3A) that play key roles in the metabolism of naturally occurring toxic dietary compounds. We found that woodrats show expansions of all three p450 gene families, including the evolution of multiple novel gene islands within the 2B and 3A subfamilies. Our assemblies demonstrate that trio binning from an F_1_ hybrid rodent effectively recovers parental genomes from species that diverged more than a million years ago. Turnover and novelty in detoxification gene islands in herbivores is widespread within distinct p450 subfamilies, and may have provided the crucial substrate for dietary adaptation during environmental change.

## Introduction

Climate change is one of the largest threats currently facing life on Earth. Under optimistic projections, a mean rise of 2°C over pre-industrial temperatures is expected by the end of the 21^st^ century (Raftery et al. 2017), and land areas are projected to experience even more severe increases (Intergovernmental Panel on Climate Change 2018). Though these changes will affect virtually every biome in substantial ways over the course of just a few decades, the potential consequences this will have on ecosystems are just beginning to be understood. One result that does seem likely, based on analyses of small climate change events over the past few centuries, are vast alterations to vegetation (Rosenzweig et al. 2007). The effects this will have on the animals that rely on these plants for sustenance – and vertebrate herbivores in particular – are not entirely known, but significant changes in the nutritional profile of available plants are virtually assured. By studying woodrats of the genus *Neotoma*, which have survived just such an event by adapting to a novel and toxic food resource, we can gain insight into the physiological and genetic factors involved in response to climatic changes and overcoming substantial dietary shifts.

Woodrats (*Neotoma* spp.) comprise the sister genus to *Peromyscus* deer mice (Figure 1A), and are endemic to North and Central America (Hall 1982). Though most woodrat species have fairly distinct ranges across the continent, there are a number of known areas where two closely related species come into contact, and reproductively viable individuals can arise when interbreeding occurs in these hybrid zones (Patton 2007; Shurtliff et al. 2014; Coyner et al. 2015). Around 17,000 years ago, the southwestern United States, which is home to several *Neotoma* species, experienced a temperature increase as the Late Glacial Cold Stage came to an end. This shift caused substantial disruption to the vegetation throughout the region and resulted in the establishment of the arid deserts of the American Southwest (Van Devender 1977; Van Devender and Spaulding 1979; Wells and Woodcock 1985). In the wake of this climate change event, juniper (*Juniperus spp.*) (Figure 1B) populations declined, and large portions of their former range were replaced with what ultimately became the dominant shrub in the region (and a defining character of the Mojave Desert): creosote bush (*Larrea tridentata*) (Figure 1C) (Wells and Hunziker 1976; Van Devender 1977; Van Devender and Spaulding 1979). Though the resident woodrat species were capable of ingesting a diet containing high levels of terpenes (the defensive compounds produced by juniper, one of their primary food sources), the shift to a juniper-depleted diet with high levels of creosote and its radically different phenolic secondary compounds presumably exerted a selective force on *Neotoma bryanti* (Figure 1D) and *N. lepida* (Figure 1E). The extent of any resulting dietary and metabolic adaptations to creosote consumption remains unclear, but the persistence of ancestral juniper-feeding populations of both species in areas north of the Mojave provides a unique opportunity to investigate their genetic underpinnings. By comparing and contrasting the genomes of creosote feeders to their juniper-feeding relatives, we can investigate the genomic mechanisms that enabled woodrats to rapidly adapt to a novel dietary resource and, in the process, endure a profound climatic shift.

**Figure 1:**
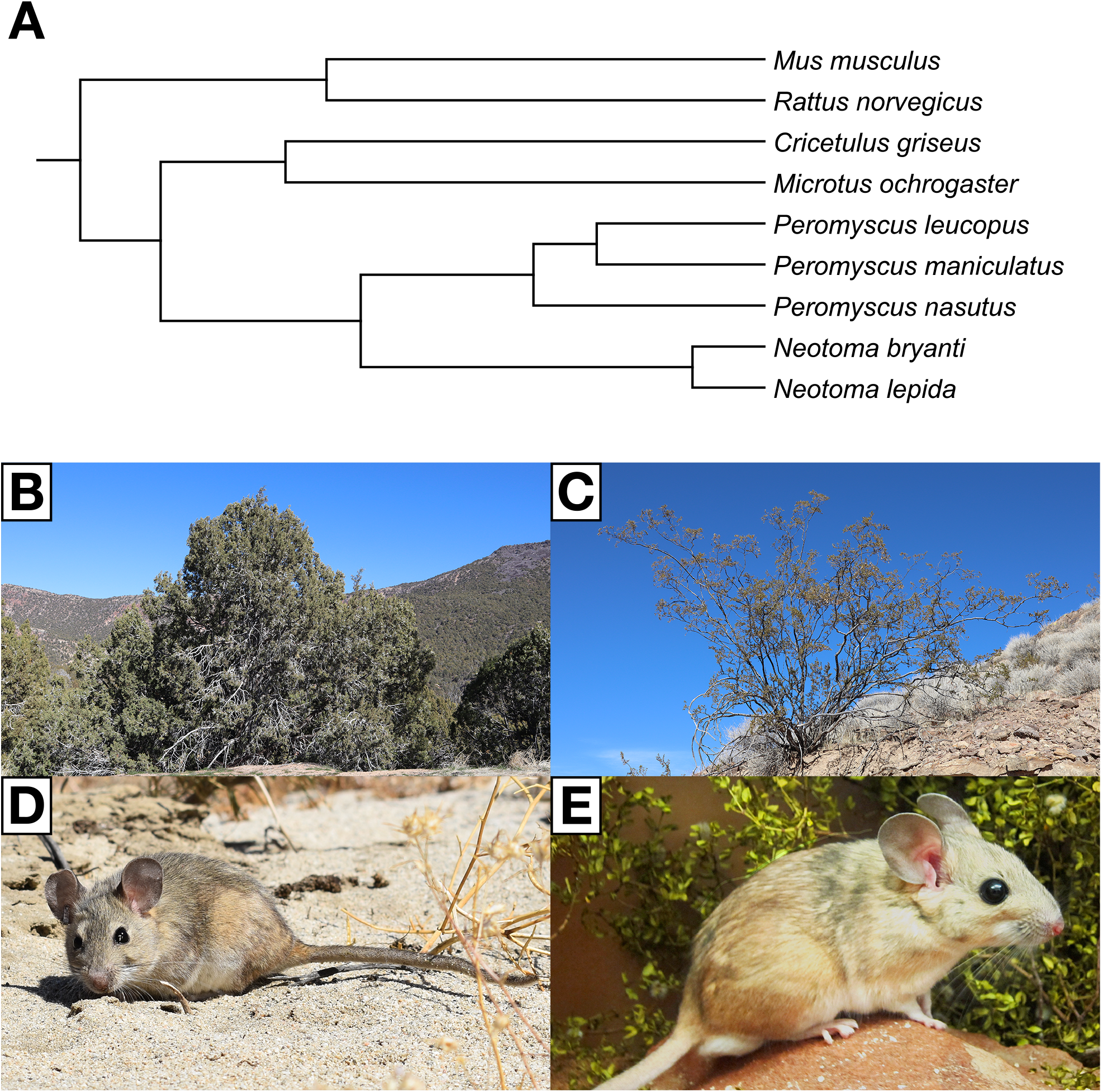
Phylogeny of Cricetidae rodents, common dietary items, and photographs of *Neotoma bryanti* and *Neotoma lepida*. (**A** a) *Neotoma* woodrats are closely related to deer mice of the genus *Peromyscus* and more distantly related to other members of the family Muridae. The phylogeny for this panel was downloaded from http://www.timetree.org and visualized in MEGA X (Stecher et al. 2020). (**B**, **C**) Images of juniper (*Juniperus spp.*) (**B**), an ancestral *Neotoma* food source, and creosote bush (*Larrea tridentata*) (**C**), the shrub that replaced it in many low elevation localities in the Mojave Desert. (**D**, **E**) Photographs of *N. bryanti* (**D**) and *N. lepida* (**E**), the two woodrat species selected for the trio binning approach.

Identifying and characterizing the putative genetic basis of these adaptations will require high-quality and annotated reference genomes, however, and until now these resources were lacking for *Neotoma* woodrats. A draft genome for *N. lepida* (Campbell et al. 2016) was the sole genomic reference assembly available for the genus, and while fairly complete, it is also highly fragmented (scaffold N50 length of 151 kb), thereby limiting our ability to perform complex genetic analyses. To address this deficiency, we leveraged the fact that the *Neotoma* species of interest are capable of hybridizing and applied the newly developed trio binning approach implemented in Canu (Koren et al. 2018) to an F_1_ cross between *N. bryanti* and *N. lepida* – sister species with an estimated 1.5-million-year divergence time (Patton 2007) – to generate new reference genomes for both organisms. By assembling each of the parental haplotypes of the F_1_ offspring separately, we were able to avoid the issues that often plague the construction of diploid vertebrate genomes while simultaneously keeping our sequencing and computational costs low. Though this technique has previously been used to generate genome assemblies for hybrid species within the Bovidae (Rice et al. 2020) and Felidae (Bredemeyer et al. 2020) families, to our knowledge this marks the first time this approach has been used to generate genomes for two rodent species, and more generally in two species of wild mammal that naturally hybridize.

The resulting references are assembled to the chromosome level, and their sequence content and annotations are highly complete by multiple metrics. In an effort to better understand the genomic basic of adaptation to different toxic diets, we interrogated the genetic architecture and possible expansion dynamics of the cytochrome p450 (henceforth p450) 2A, 2B, and 3A gene families. The p450 enzymes produced by these genes are critical in the biotransformation process of toxins like those present in creosote bush and juniper (Magnanou et al. 2009; Skopec et al. 2013). Novel dietary niches involving toxic food are hypothesized to exert strong selection pressure for herbivore defense against toxic effects. Previous work in these and other woodrats suggests gene copy number evolution has brought about both novel function and induced overexpression in dietary specialists (Magnanou et al. 2009; Malenke et al. 2012; Kitanovic et al. 2018). If gene copy number evolution is an important part of adaptation to these novel diets, then we expect to document copy number expansion of the 2B and potentially other p450 gene subfamilies in comparison to related Old and New World Rodents.

## Materials and Methods

### Animal collection and handling

All animal work was approved under the University of Utah’s Institutional Animal Care and Use Committee (Protocol 16_02011). The female *N. bryanti* used in the F_1_ cross was captured in March 2018 near Pioneertown, California, USA (lat. 34.151, lon. −116.479). The male *N. lepida*, kindly provided by Dr. James Patton, was captured in October 2018 near Kelso Depot, Mojave Preserve in California, USA (lat. 35.009, lon. −115.645). *Neotoma lepida* animals used for the transcriptomics portion of the work were captured in March 2018 in Lytle Ranch Preserve, Utah, USA (lat. 37.070, lon. −114.000). All individuals were live-trapped at their respective sites using long (7.62 × 89 × 22.86 cm) Sherman traps. The *N. lepida* captured near Kelso Depot was obtained under the California Department of Fish and Wildlife permit number SC-2105, MOJA-00038 issued to James Patton, while all other animals were trapped under the California Fish and Game permit SCP-8123 and Utah Game and Fish permit 1COLL5194-2 issued to M. Denise Dearing.

After transport to the animal facility in the School of Biological Sciences at the University of Utah, animals were housed in solid-bottom shoebox cages (48 × 27 × 20 cm) containing a plastic tube and lined with cedar shavings. The facility was maintained at 28°C, 15-20% relative humidity and with a 12-hour light/dark cycle. Animals were provided rabbit chow and water *ad libitum*. With the exception of the mating required for the F_1_ cross (and the resulting pups), animals were housed individually. The *N. bryanti* × *N. lepida* cross yielded three hybrid individuals (two females, one male) in March 2019, and one female from this litter was selected for sequencing to ensure the X chromosome could be reconstructed for each haplotype.

All seven animals (one *N. bryanti*, five *N. lepida*, one hybrid) were dispatched in accordance with IACUC protocols prior to dissection. For DNA sequencing, liver tissue was extracted from each individual and stored at −80°C. For RNA sequencing, extracted liver tissue was first placed in RNAlater (ThermoFisher Scientific, Waltham, Massachusetts, USA) prior to storage at −80°C.

### Genomic library preparation and sequencing

For the *N. bryanti* and *N. lepida* parents, DNA was extracted from liver tissue using the DNeasy Blood & Tissue Kit (QIAGEN, Hilden, Germany), and the concentration (approximately 70 ng/μL) was validated using a Qubit dsDNA HS Assay Kit (ThermoFisher Scientific, Waltham, Massachusetts, USA). Following shearing with a Covaris S2 Focused-ultrasonicator (Covaris, Woburn, Massachusetts, USA), libraries with an average insert size of 450 bp were prepared by the University of Utah’s Huntsman Cancer Institute High-Throughput Genomics Core Facility using the Illumina TruSeq DNA PCR-Free Library Prep Kit (Illumina, San Diego, California, USA). 2 × 150 bp reads for each parent were generated on an Illumina NovaSeq 6000 instrument using the NovaSeq S2 reagent kit (Illumina, San Diego, California, USA). A total of 539,570,835 reads for the *N. bryanti* mother and 648,777,162 reads for the *N. lepida* father were produced, yielding 67.44× and 81.10× coverage, respectively, based on a genome size estimate of 2.4 Gb (this estimate was derived from the *N. lepida* ASM167557v1 draft genome assembly [Campbell et al. 2016], which, while fragmented, was reported to be quite complete).

For the hybrid pup, DNA extraction, library preparation and sequencing were carried out by the DNA Sequencing Center at Brigham Young University. Following DNA extraction with the QIAGEN MagAttract HMW DNA Kit (QIAGEN, Hilden, Germany), libraries were generated using the SMRT Bell Template Preparation Kit (Illumina, San Diego, California, USA), and sequencing was performed using eight SMRT Cells on a PacBio Sequel II instrument (Illumina, San Diego, California, USA). This yielded 787.76 Gb of sequence data, corresponding to approximately 328× coverage based on the 2.4 Gb genome size estimate.

### RNA library preparation and sequencing

RNA was extracted from liver tissue using the QIAGEN RNeasy Mini Kit (QIAGEN, Hilden, Germany). The concentration (approximately 75 ng/μL) and RNA quality (RIN ≥ 9.0) for each sample was validated with a Qubit RNA BR Assay Kit (ThermoFisher Scientific, Waltham, Massachusetts, USA) and Agilent Technologies RNA ScreenTape Assay (Santa Clara, California, USA), respectively, before library preparation and sequencing was carried out by the University of Utah’s Huntsman Cancer Institute High-Throughput Genomics Core Facility. Libraries were generated using the Illumina TruSeq Stranded mRNA Library Prep Kit (Illumina, San Diego, California, USA), and 2 × 150 bp reads were produced on an Illumina NovaSeq 6000 instrument using the NovaSeq S2 reagent kit (Illumina, San Diego, California, USA). An average of 57,521,215 reads were generated for each of the 4 samples. For exact counts, see Table S1.

### Mitochondrial genome assembly

Illumina short reads for each parent were imported into CLC Genomics Workbench 12 (https://digitalinsights.qiagen.com), trimmed using the default settings of the “Trim Reads” tool and assembled using the default settings of the “De novo assembly” tool. For each of these assemblies, the sequences were aligned with BLAT 36 (Kent 2002) against the *Rattus norvegicus* Rnor 6.0 mitochondrial genome (Rat Genome Sequencing Project Consortium 2004) to identify which scaffolds corresponded to the mitochondria. For each parent, the mitochondrial genome appeared to be contained in a single sequence (17365X2_contig_1766 for *N. bryanti* and 17365X1_contig_1575 for *N. lepida*), and subsequent BLASTN (Camacho et al. 2009) searches against the online NCBI NR database (O’Leary et al. 2016) on December 3, 2020 confirmed these findings.

### Read partitioning and haplotype assembly

PacBio reads were converted from the BAM to the FASTA format using version 1.3.0 of the bam2fasta tool (https://pacb.com) for assembly. Employing *k*-mers identified in the Illumina read data generated from the parents, Canu 1.9 (Koren et al. 2018) was used to partition the PacBio reads of the hybrid by haplotype (and therefore species). The resulting species-specific reads were assembled separately using the default settings of Canu with an estimated genome size of 2.4 Gb.

### Genome polishing

BAM files containing species-specific reads were generated from the original PacBio subread BAMs using SAMtools 1.9 (Li et al. 2009) and a custom Python script that parsed read names from the Canu haplotype-specific FASTAs; these BAM files were subsequently indexed using pbindex 1.0.7 (https://pacb.com) to generate input files compatible with the pbmm2 aligner. Using the default settings of pbmm2 1.2.1 (https://pacb.com; Li 2018), these reads were aligned against the appropriate Canu reference for each species. Arrow 2.3.3 (https://pacb.com) was run with default settings to polish each assembly; due to the high memory requirements of this program, the pbmm2 BAMs and Canu reference FASTAs were split into individual files by contig using SAMtools 1.9 and custom Python scripts employing the Biopython 1.76 libraries (Chapman and Chang 2000). Arrow was run on each contig individually, and the resulting FASTA sequences were joined once the polishing was complete. These polished assemblies were subsequently subjected to a second round of pbmm2 alignment and Arrow polishing to generate the final contig sequences.

### Identification and masking of repetitive elements

RepeatModeler 2.0.1 (Flynn et al. 2020) was run with the LTR structural search pipeline enabled on each polished assembly to generate species-specific repeat libraries. Utilizing these species-specific libraries as well as the 20181026 version of the RepeatMasker libraries from Repbase (Jurka 1998), RepeatMasker 4.0.9 (http://www.repeatmasker.org) was used to mask repetitive elements in each assembly. As the resulting masked FASTAs were to be used for automated gene annotation, RepeatMasker was run with the “-nolow” argument to avoid masking low complexity regions that could potentially comprise parts of genes. RepeatMasker results for each genome are contained in Table S2 and S3.

### Processing and alignment of transcriptomic reads

The transcriptomic reads generated from this study were supplemented with the RNA-seq reads used to annotate the previous ASM167557v1 draft release of the *N. lepida* genome (Campbell et al. 2016). Quality control and trimming of all RNA-seq reads was performed with Trim Galore! 0.65 (https://www.bioinformatics.babraham.ac.uk/projects/trim_galore/), which utilized FastQC 0.11.9 (http://www.bioinformatics.babraham.ac.uk/projects/fastqc/) and Cutadapt 2.10 (Martin 2011). Out of the initial 313,541,822 reads, 312,133,692 (99.55%) were retained following Trim Galore! processing. Trimmed reads were subsequently aligned to each genome using the two-pass mode of STAR 2.5.7a (Dobin et al. 2013), and merged into a single BAM file with SAMtools 1.9 (Li et al. 2009). Mapping statistics for each RNA-seq sample are available in Table S1.

### Automated genome prediction

Gene models on the repeat-masked references were predicted with BRAKER 2.1.5 (Hoff et al. 2019). The STAR RNA-seq alignments, as well as all protein sequences from the OrthoDB v10 database (Kriventseva et al. 2019), were supplied to the BRAKER command line arguments “--bam” and “--prot_seq”, respectively, and the application was run using the “--etpmode” and “--softmasking” arguments. Gene hints from the protein sequences and RNA-seq alignments were identified by the GeneMark-ES 4.59 suite (Brůna et al. 2020), and a subset of these predictions determined to be high quality based on the supporting OrthoDB and transcriptome evidence were selected by BRAKER for training. After aligning these predictions against each other with DIAMOND 2.0.0 (Buchfink et al. 2015) to remove redundant models, these high-quality genes were used to train AUGUSTUS 3.3.3 (Stanke et al. 2004) parameters for each species prior to gene prediction. Custom Python scripts were used to apply minor reformatting to the resulting BRAKER GFF3 so that it could be sorted and validated by GenomeTools 1.6.1 (Gremme et al. 2013).

### Gene model filtering and functional annotation

Protein sequences for all predicted transcripts were obtained from each species’ BRAKER GFF3 and polished reference FASTA with Python scripts utilizing the Biopython 1.76 libraries (Chapman and Chang 2000). InterProScan 5.32-71.0 (Jones et al. 2014), was run on each of these sequences, and all gene predictions lacking an InterPro group ID were removed from the BRAKER annotations. The resulting GFF3s were then analyzed and processed by version 2.0.1 of the GFF3toolkit (https://github.com/NAL-i5K/GFF3toolkit) to resolve incomplete and spuriously duplicated gene models. To provide additional functional annotation, all remaining protein sequences were aligned with BLASTP 2.9.0+ (Camacho et al. 2009) against a database containing protein sequences from *Homo sapiens* (release GRCh38 [Schneider et al. 2017]), *Mus musculus* (release GRCm38.p6 [Mouse Genome Sequencing Consortium 2002]), *Peromyscus leucopus* (release Pero 0.1 [Long et al. 2019]), *Peromyscus maniculatus* (release Pman 2.1, downloaded from Ensembl [Yates et al. 2019]) and *Rattus norvegicus* (release Rnor 6.0 [Rat Genome Sequencing Project Consortium 2004]). When a protein prediction had a best hit with an E-value exceeding 10^−5^, a description and gene symbol (if available) were retrieved from the corresponding genome annotation and added to the BRAKER GFF3 alongside all relevant InterPro group IDs.

### Annotation of p450 genes

We leveraged several sources of additional information to identify and then manually curate the p450 genes in each woodrat genome. We produced a set of p450 gene models in BITACORA v1.3 (Vizueta et al. 2020), which combines Hidden Markov Model (HMM) protein profiles and contig aware linear searches to better annotate tandemly arrayed genes. We employed the default BITACORA settings and the Pfam PF00067 HMM profile for the p450s in these searches. Additionally, we used the p450 amino acid sequence dataset from Thomas (Thomas 2007) to query our genomes using TBLASTN searches (version 2.11, e-value = 0.00001). Finally, we used HISAT2 v.2.2.1 (Kim et al. 2019) to conduct splice aware RNA-seq alignments to each genome using the “--no-unal --no-mixed --no-discordant” options, and filtered the resulting alignment map files in SAMtools 1.9 (Li et al. 2009) to retain only the top alignment for each read pair. An additional GFF3 was also generated by aligning CDS sequences from the five animal genomes employed for functional annotation to each of the woodrat references using GMAP version 2020-06-01 (Wu and Watanabe 2005) with “--min-trimmed-coverage” set to 0.8. The resulting GFF3 was processed with version 2.0.1 of the GFF3toolkit (https://github.com/NAL-i5K/GFF3toolkit) and sorted and validated with GenomeTools 1.6.1 (Gremme et al. 2013).

### Manual curation of cytochrome P450s

The contigs, p450 blast hits, RNA-seq alignments, and various gene model annotations were loaded into Geneious 10.2.6 (https://www.geneious.com) for manual curation of p450 genes. The p450 gene models from the combined BRAKER, GMAP, and BITACORA annotations were used as the basis for producing a final quality model with the following characteristics: AG/GT splice sites, appropriate reading frames, and a lack of internal stop codons. When RNA-seq data covered the entire model, the splice sites supported by the RNA-seq alignments were given priority. Putative p450s with early stops, or those with largely incomplete coding sequences based on TBLASTN searches of the annotated product were marked as pseudogenes. Pseudogene designations included any p450 blast hit that was near a functional gene or p450 gene island, and ranged from essentially complete sequences to hits matching single exons. We then exported curated Geneious GFF3 annotations for each p450 gene.

### Inference of syntenic p450 gene islands and copy number differences

We visualized the genomic organization of p450 2ABFGST and 3A genes for both *Neotoma* as well as an Old World rat (*R. norvegicus*) and New World mice (*P. maniculatus* for the cyp2ABFGST genes and *P. leucopus* for the cyp3A genes). These two different *Peromyscus* were chosen based on the apparent completeness of the gene clusters in each species, so inference of expansions in *Neotoma* represents the most conservative comparison that could be made to its sister genus based on available genomic resources. We also visualized nearby non-p450 genes to help establish regional synteny. We accomplished these visualizations using the Geneious 10.2.6 viewer. The amino acid sequences of all functional genes within each subfamily were exported from Geneious and aligned in MAFFT 7.45 (Katoh and Standley 2013) using the G-INS-I algorithm and the BLOSUM62 scoring matrix. We then used the “backtrans” function in TreeBeST 1.9.2 (http://treesoft.sourceforge.net) to backtranslate the amino acid alignment to a protein-guided codon alignment of nucleotides sequences, and next used the TreeBeST “best” function to derive gene trees guided by the species tree with nodal support derived from 100 bootstraps. The species tree used was downloaded from www.timetree.org. The models of each gene island and gene trees were then edited in Adobe Illustrator (Adobe Inc., San Jose, California, USA), using both gene order and the clustering in the gene tree to guide the calling of putatively syntenic p450 gene clusters.

### Resolution of the genomic region containing the cyp2B array in *N. lepida*

While the cyp2B array in *N. bryanti* was assembled as a single contig in the Canu assembly, the *N. lepida* array was significantly more fragmented, with complete, partial and pseudogenized cyp2B sequences identified across seven contigs (Nlep_tig00001124, Nlep_tig00001166, Nlep_tig00001922, Nlep_tig00004740, Nlep_tig00010068, Nlep_tig00418826 and Nlep_tig00418827). To resolve the region in *N. lepida*, haplotype-specific PacBio reads were aligned to the *N. lepida* assembly with minimap2 2.17-r941 (Li 2018) and only those mapping to the seven identified contigs were retained. Two Canu 1.9 (Koren et al. 2018) assemblies were constructed with these reads: the first was generated using all reads (hereafter “AR”; genome size was left at 2.4 Gb and stopOnLowCoverage was set to 0.7), while the second was generated using Canu’s default coverage settings (hereafter “40×”; genome size was set to 19 Mb, which was the approximate size of the seven contigs containing cyp sequences). Each of these assemblies was subsequently polished using two rounds of pbmm2 1.2.1 (https://pacb.com; Li 2018) and Arrow 2.3.3 (https://pacb.com).

The 40× assembly appeared to resolve virtually all of the region into two contigs: one 6.58 Mb and another 11.98 Mb in length. Though more fragmented overall, the AR assembly contained a single contig 13.63 Mb in length that, based on MUMmer 3.23 (Kurtz et al. 2004) alignments (Figure S10A, B), contained portions of both contigs from the 40× assembly. Using quickmerge 0.3 (Chakraborty et al. 2016) with --length_cutoff set to 1,000,000, the two 40× contigs were aligned against the 13.63 Mb AR contig to generate a single sequence 18.55 Mb in length. To polish this sequence, reads were first aligned with minimap2 to the AR and 40× assemblies and only those mapping to the three contigs of interest were retained; these reads were then used for two rounds of polishing with pbmm2 and Arrow. Based on MUMmer alignments, the resulting sequence resolved the order and orientation of the largest four contigs present in the initial assembly (Figure S10C). The remaining three contigs could not be definitively placed and appeared to be repetitive sequences possibly arising from unresolved sequencing errors (Figure S10D). RepeatMasker 4.0.9 (http://www.repeatmasker.org) and the *N. lepida* RepeatModeler 2.0.1 (Flynn et al. 2020) library were used to mask repetitive elements on this sequence as previously described, and gene models from the initial Canu assembly were transferred to this sequence using Geneious 10.2.6 (https://www.geneious.com).

### Assessing genome completeness

Genome completeness for both *Neotoma* genomes was assessed using the “genome” mode of BUSCO against the Glires_odb10 dataset of 13,798 genes. We ran both BUSCO 4.1.4 (Seppey et al. 2019), which employed gene predictions of both AUGUSTUS and TBLASTN, and the newer BUSCO version 5, which employed the metaeuk gene search for eukaryotic genomes. The results of these searches were largely overlapping, but some genes were found by only one version. We combined the gene lists from each run to produce our final BUSCO completeness metrics (Table 1).

**Table 1:**
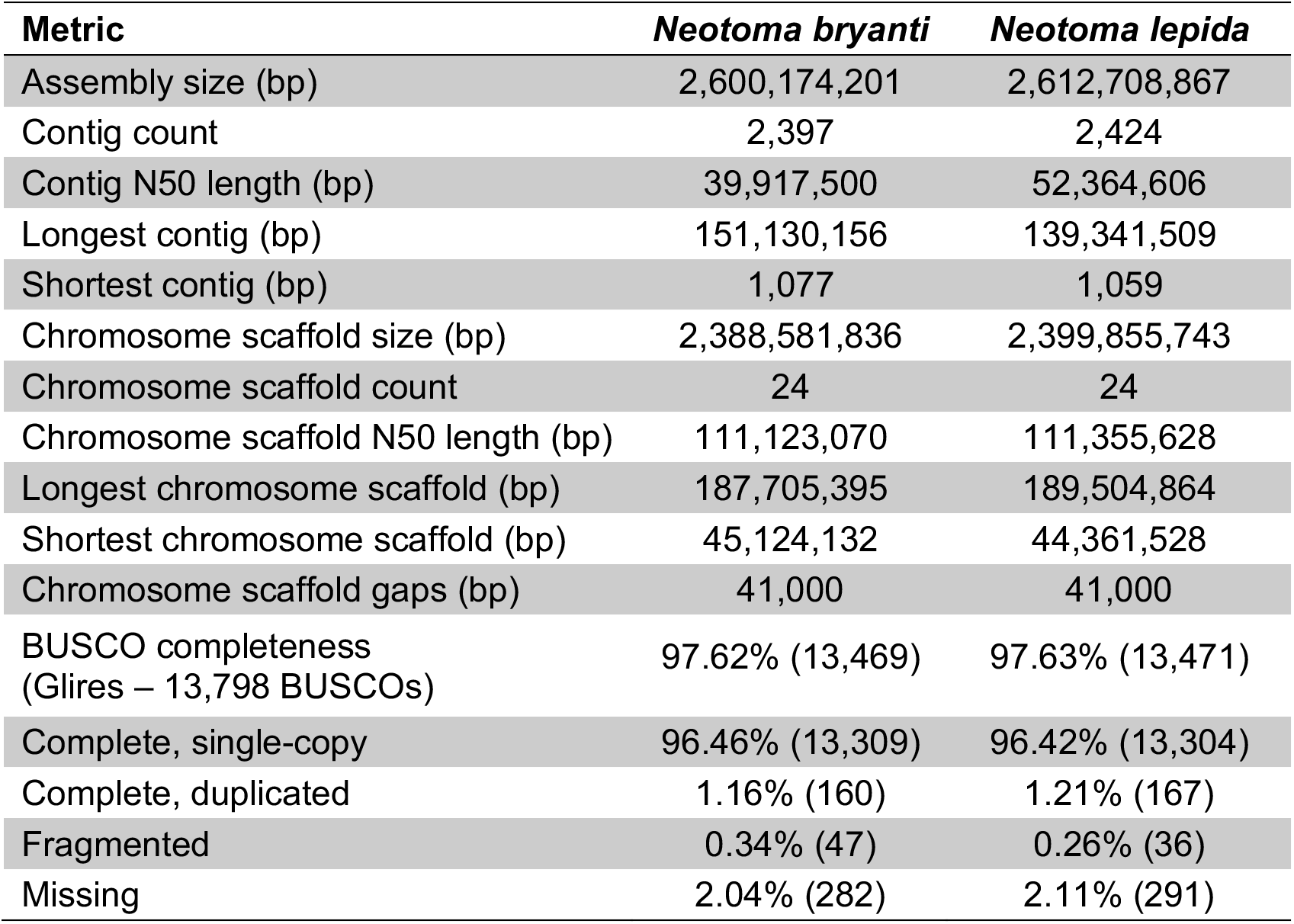
Assembly metrics and BUSCO assessments for *Neotoma bryanti* and *Neotoma lepida*.

Jellyfish 2.3.0 (Marçais and Kingsford 2011) was used to tabulate *k*-mer counts of length 25 in the trimmed and corrected reads generated for each of the *Neotoma* assemblies. After discarding low count *k*-mers presumed to be noise (those that appeared less than 8 times for *N. bryanti* and 11 times for *N. lepida*), the genome size was estimated by multiplying each count value by the total number of *k*-mers present at that value, then dividing that number by the *k*-mer count value that appeared most frequently (24 for *N. bryanti* and 30 for *N. lepida*). Plots of the *k*-mer distributions for *N. bryanti* and *N. lepida* are available in Figure S1.

### Genome scaffolding and syntenic analysis

Contigs for both woodrat species were aligned to the chromosome scaffolds from the reference assemblies of *Peromyscus leucopus* (white-footed mouse, release PerLeu 2.1 [Long et al. 2019])*, P. maniculatus* (deer mouse, release Pman 2.1, downloaded from Ensembl [Yates et al. 2019]) and *P. nasutus* (northern rock mouse, available from https://www.dnazoo.org/assemblies/Peromyscus_nasutus/), the closest relatives for which chromosome-level assemblies were available. Chromosome numbering and orientation are based on the *P. maniculatus* reference, with the corresponding chromosome sequences in *P. leucopus* and *P. nasutus* determined by alignments of those assemblies against the *P. maniculatus* reference using the default settings of MUMmer 3.23 (Kurtz et al. 2004). Due to the potential for misalignment due to repetitive elements, only *Neotoma* contigs exceeding 1 Mb in length with at least 100 kb of sequence uniquely aligned to a single chromosome were used in the scaffolding process. Contigs were aligned with the default settings of MUMmer 4.0.0 beta 2 (Marçais et al. 2018), and assigned to the chromosome they had the greatest amount of sequence uniquely aligned to. Gaps between adjacent contigs were denoted by a stretch of 500 N characters in each assembly. The orders and orientations of contigs along each chromosome were determined by the median position of their alignments, and were manually adjusted if discontinuities were evident when the MUMmer alignments were visualized (these discontinuities were almost universally large-scale inversions relative to the *Peromyscus* reference sequences). Once preliminary chromosome scaffolds were generated against the *Peromyscus* genomes, the *N. bryanti* and *N. lepida* sequences were aligned against each other for validation as well as to correct any outstanding contig placement issues.

After scaffolding, syntenic regions between each *Neotoma* genome and *P. leucopus* and *P. maniculatus* were identified using BLASTALL 2.2.26 (Altschul et al. 1990) protein alignments with an E-value of 10^−20^ and the default settings of the latest Synima GitHub release (a0bc445, dated June 2, 2020) (Farrer 2017). MUMmer plots generated with a minimum cluster size of 250 against the *Peromyscus* genomes are available for all *N. bryanti* chromosomes as Figures S2-S4, and all *N. lepida* chromosomes as S5-S7. MUMmer plots generated with a minimum cluster size of 2,500 between the *Neotoma* chromosome scaffolds are available in Figures S8 and S9. Information on the relative order and orientation of contigs for each chromosome is contained in Table S4.

### Estimating nuclear and mitochondrial divergence

The *N. bryanti* and *N. lepida* genome assemblies were aligned to each other using the NUCmer program of MUMmer 3.23 (Kurtz et al. 2004), and delta-filter was run on the subsequent alignments to retain only regions that had one-to-one mappings between both species. Variants in these regions were then called using the MUMmer show-snps program, and metrics were calculated using a custom Python script. The same process was performed for the mitochondrial scaffolds. Complete statistics for both the nuclear and mitochondrial genomes are contained in Table S5.

## Results and Discussion

### Read binning produces highly contiguous and complete assemblies for hybridizing rodent species

The read lengths provided by recent advances in sequencing technology, combined with the trio binning approach, yielded an almost perfect separation of sequencing data from the F_1_ offspring. After removing all reads considered too short (those less than 1 kb in length, which accounted for 1.97 Gb or 0.25% of the total), Canu assigned 49.46% (389.60 Gb) of reads to the maternal *N. bryanti* haplotype and 50.16% (395.14 Gb) to the paternal *N. lepida* haplotype. Only 0.13% (1.05 Gb) was unassignable. This marks a slight advancement over the results reported for trio binning within the *Bos taurus* species (49.3% and 49.6% assigned to each haplotype, respectively) (Koren et al. 2018), and is in line with the results reported for the more recent cross-species *Bos* (Rice et al. 2020) and feline (Bredemeyer et al. 2020) assemblies. The fact that the highly accurate partitioning of trio binning yielded minimal loss of usable sequencing data further demonstrates the promise this technique holds for generating genomes from hybridizing species.

Though well over 2,000 contigs were produced for each woodrat reference, the N50 values indicate that the vast majority of each assembly lies within large, contiguous sequences (Table 1). Furthermore, although fewer than 5% of the contigs in each assembly exceeded 1 Mb in length (110 for *N. bryanti* and 109 for *N. lepida*), these sequences contained over 91% of each 2.6-Gb assembly. Thus, while small contigs are numerous, they make up only a small portion of each reference genome. Of particular note were the handful of extremely long contigs generated for each assembly; with lengths exceeding 100 Mb, each of these sequences likely represent most (or all) of a chromosome.

Analysis of the assemblies with BUSCO indicated that the genomes were remarkably complete, with *N. bryanti* containing 97.62% and *N. lepida* possessing 97.63% of the 13,798 single-copy orthologs present in the Glires version 10 dataset (Table 1). Additionally, both assemblies contained few partial or multi-copy BUSCOs, indicating that fragmentation and sequence duplication were kept to a minimum. The total assembly lengths were also similar to genome size estimates based on *k*-mer counts from the PacBio sequencing data (2.72 Gb and 2.62 Gb for *N. bryanti* and *N. lepida*, respectively), with the assembled and estimated values differing by less than 5%. Taken together, these results indicate that the vast majority of the nuclear genome for each species is contained in the assemblies.

Similar results were found for the mitochondrial genomes reconstructed from the parental short read data. Each mitochondrial scaffold – 16.7 kb for *N. bryanti* and 16.3 kb for *N. lepida* – was very close in size to the 16.3-kb mitochondrial assemblies for *R. norvegicus* (Rat Genome Sequencing Project Consortium 2004) and *M. musculus* (Mouse Genome Sequencing Consortium 2002), indicating that the *Neotoma* scaffolds contain virtually all of the expected mitochondrial sequence. These findings were strengthened with subsequent BLASTN (Camacho et al. 2009) searches against the online NCBI NR database (O’Leary et al. 2016), which found query coverages of 97% and 99% for *N. bryanti* and *N. lepida*, respectively, for the closely related *N. magister* and *N. mexicana* mitochondrial genomes. Much like the nuclear genomes, these results demonstrate that the *N. bryanti* and *N. lepida* mitochondrial genomes are complete.

### Scaffolding indicates most chromosomes were assembled from few contigs

Using the 24 chromosome sequences from each of the *Peromyscus leucopus*, *P. maniculatus* and *P nasutus* genomes as references, we were able to scaffold 96.36% of contigs exceeding 1 Mb in length for *N. bryanti* (106/110), and 99.08% (108/109) for *N. lepida* (Figure 2A, B). The scaffolded chromosomes resulting from this approach contained the vast majority of each species’ genome, comprising 91.86% and 91.85% of the total *N. bryanti* and *N. lepida* assemblies, respectively (Table 1). Though the number of chromosomes in *Neotoma* (1N = 26 chromosomes in *N. lepida*) may be slightly higher than in *Peromyscus* (Mascarello and Hsu 1976), given the close phylogenetic proximity of these genera, our scaffolding approach should provide a reasonably accurate representation of each *Neotoma* species’ chromosomal architecture.

**Figure 2:**
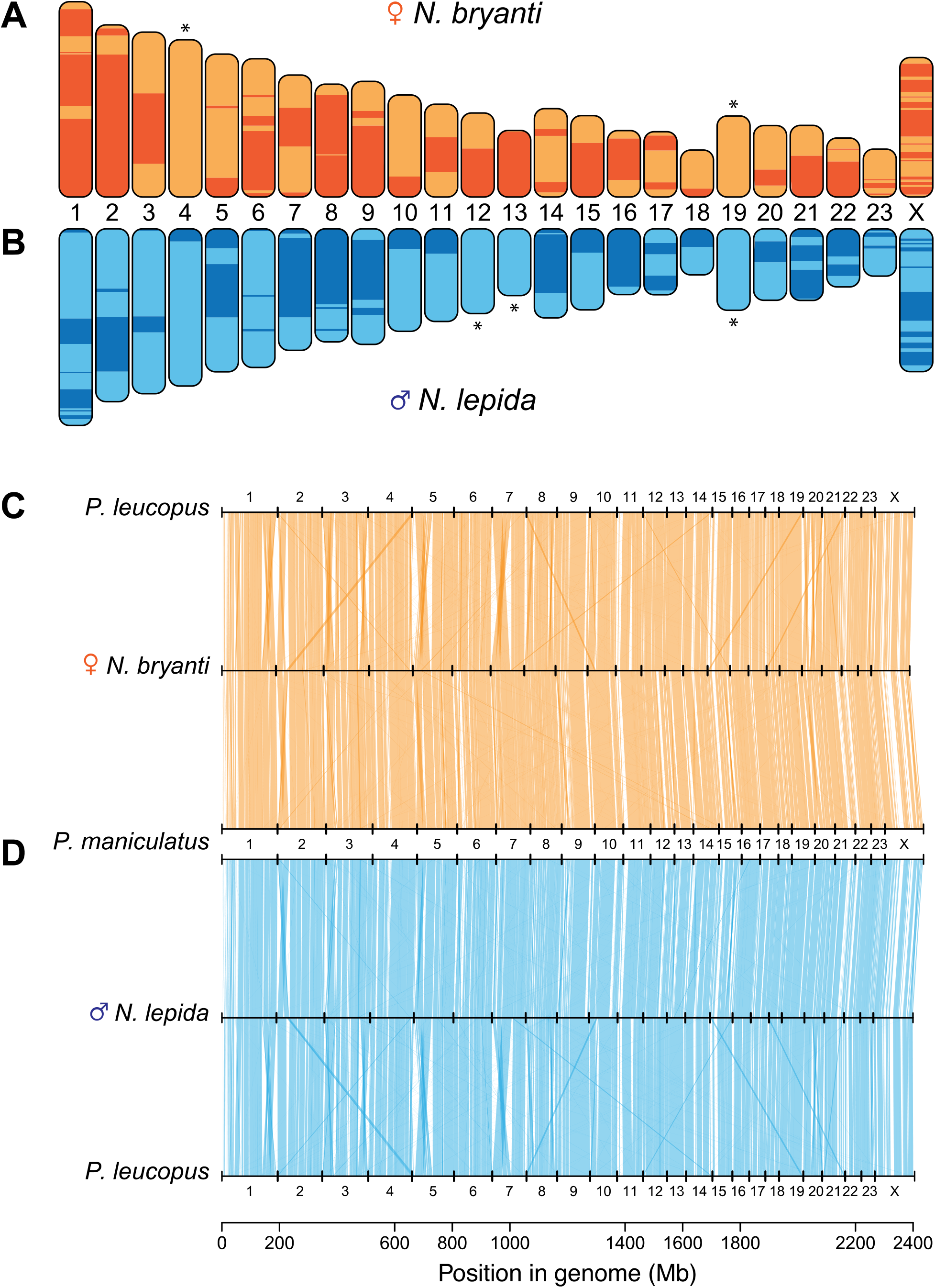
Contig scaffolding and synteny results for both *Neotoma* genomes. (**A**, **B**) Visualization of contigs scaffolded into chromosomes for *Neotoma bryanti* (**A**) and *N. lepida* (**B**). Alternating colors denote individual contigs, and chromosomes indicated with an asterisk were assembled in their entirety as a single contig. (**C**, **D**) Identification of syntenic gene alignments by Synima for the *N. bryanti* (**C**) and *N. lepida* (**D**) autosome scaffolds 1-23 and X chromosome scaffold against the *Peromyscus leucopus* and *P. maniculatus* genomes. Note that while there are a number of instances of cross-alignments among the chromosomes (likely due to cross-mapping between genes in duplicated families), the positions of the majority of genes are strongly conserved between *Neotoma* and *Peromyscus*.

Among the assembled chromosomes, five were found to be assembled as single contigs; these were chromosomes 4 and 19 for *N. bryanti* and 12, 13 and 19 for *N. lepida* (Figure 2A, B). Further, of the chromosomes scaffolded for both species, the vast majority (23/24 for *N. bryanti* and 22/24 for *N. lepida*) were assembled from eight or fewer contigs – only the X chromosome for *N. bryanti* (23 contigs) and chromosomes 1 and X for *N. lepida* (12 and 17 contigs, respectively) were composed of more contigs (Table S4). Given that repetitive elements comprise large portions of the X chromosome in mammals (Ross et al. 2005) and can be notoriously difficult to assemble, the fragmentation of this chromosome in both assemblies is not surprising. Nevertheless, the selection of a female hybrid offspring for sequencing enabled us to reconstruct the X chromosome for both woodrat species. Along with the BUSCO and *k*-mer analyses, these scaffolding results provide further validation for the accuracy and completeness of the nuclear genome assemblies.

### Large-scale chromosome structure and synteny are conserved between *Neotoma* and *Peromyscus*

Alignments of the scaffolded chromosome sequences of the two *Neotoma* species to the three available *Peromyscus* genomes reveal that the majority of the chromosomal sequence is present in the same order and orientation in both genera of rodents (Figures S2-S7). While a number of structural rearrangements (primarily inversions) are present, we see no evidence to suggest translocation events between chromosomes. Furthermore, the number of inversions we detected might be inflated. Very few of these structural features are present in all three *Peromyscus* references we examined (Figures S2-S7), leading to the possibility that some inferred inversions might be assembly artifacts resulting from the different scaffolding approaches used for each *Peromyscus* genome. Synteny findings, too, are broadly consistent with the alignment results, with gene positions strongly conserved between the *Neotoma, P. leucopus*, and *P. maniculatus* genomes (Figure 2C, D).

Though our alignment-based scaffolding approach cannot rule out large-scale structural differences, individual contig alignments between *N. bryanti* and *N. lepida* lend little support to their existence (Figures S8, S9). This is mildly surprising given the chromosomal variation previously observed within *Neotoma* and even within the *N. lepida* species itself (Mascarello and Hsu 1976). The lack of such structural changes, however, may prove instrumental in maintaining the viability of offspring produced when these species hybridize in the wild.

### Nuclear and mitochondrial sequence divergence between *N. bryanti* and *N. lepida*

Sequence divergence was found to be 1.27% between the *N. bryanti* and *N. lepida* nuclear genomes (Table S5), which is in line with an estimated 1.5-million-year divergence (Patton 2007). For the mitochondrial genome, divergence was 6.96% for the 16,278 bp of sequence that could be aligned between the two species. As expected, this mitochondrion-wide estimate is lower than the 9% divergence of cytochrome *b* (Patton 2007). Elevated rates of mitochondrial to nuclear divergence are common among vertebrates (Allio et al. 2017), however, and the fact that both *Neotoma* species can hybridize and produce reproductively viable offspring suggests that gene flow occurs frequently enough between the two to limit nuclear divergence.

### Gene counts and transposable element annotations are similar to other rodents

After filtering the BRAKER models based on InterPro group assignments, 22,471 protein coding genes containing a total of 26,334 transcripts were predicted for *N. bryanti*, while for *N. lepida*, 21,448 genes with 25,391 transcripts were identified. These numbers are in line with the 22,549 and 21,587 protein coding genes annotated in the current *M. musculus* (Mouse Genome Sequencing Consortium 2002) and *R. norvegicus* (Rat Genome Sequencing Project Consortium 2004) annotations, respectively. Results from RepeatMasker analysis of each genome are also broadly in agreement with predictions for the *M. musculus* (http://www.repeatmasker.org/species/mm.html) and *R. norvegicus* (http://www.repeatmasker.org/species/rn.html) genomes, with 40.62% and 41.44% of the *N. bryanti* and *N. lepida* assemblies, respectively, composed of repetitive elements (see Tables S2 and S3 for a detailed breakdown of transposable element families in each genome). Together, these annotation results for *Neotoma* are similar in overall structure to the high-quality annotations available for model rodent species.

### Dynamic evolution of cytochrome P450 gene islands and the genomic basis of toxin tolerance

After confirming the completeness of the assemblies and annotations for *N. bryanti* and *N. lepida*, we next examined the p450 family of genes due to their crucial roles in the metabolism of dietary toxins. Both p450 gene subfamilies showed clear signs of dynamic evolution and lineage-specific expansion in *Neotoma*. For the 2ABGFST genes, both *Neotoma* genomes revealed multiple instances of novel gene expansion via duplication. For example, both *R. norvegicus* and *P. leucopus* have 3 *cyp2A* genes each, while that number was expanded to 7 and 6 in *N. bryanti* and *N. lepida*, respectively (Table 2, Figure 3A). Based on the gene tree (Figure 3C), separate expansions of *cyp2A2*-like and *cyp2A3*-like genes have produced these increases in *cyp2A* total counts, and based on the relationships among species, 2A expansions in *Neotoma* rather than losses in the other genera are the most parsimonious explanation.

**Table 2:** P450 subfamily gene counts.

| **P450 Subfamily** | ***Rattus norvegicus*** | ***Peromyscus sp.*** | ***Neotoma bryanti*** | ***Neotoma lepida*** |
| --- | --- | --- | --- | --- |
| 2A | 3 | 3 | 7 | 6 |
| 2B | 8 | 9<sup>a</sup> | 7 | 15 |
| 3A | 6 | 13<sup>b</sup> | 18 | 15 |

a. *Peromyscus maniculatus* used for gene counts. b. *Peromyscus leucopus* used for gene counts.

**Figure 3:**
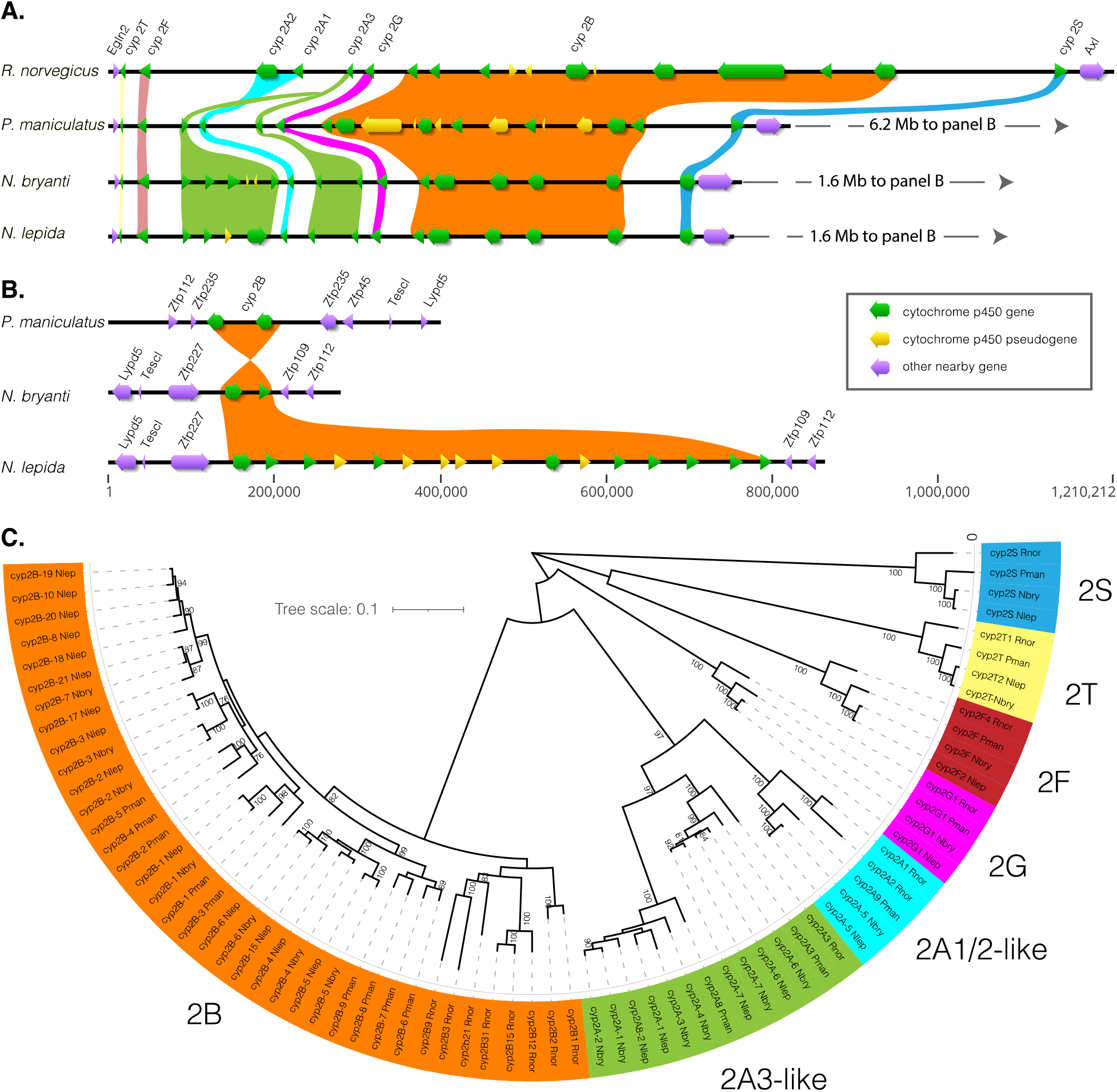
Genomic architecture and evolution of the cyp2ABFGST gene islands in four rodents. Visualization of the contigs containing these genes was based on annotations for the Norway rat (*Rattus norvegicus*; mRatBN7.2) and deer mouse (*Peromyscus maniculatus*; HuPman2.1), and manual annotation of these regions for *Neotoma bryanti* and *N. lepida*. (**A**) A highly conserved gene island is bounded by *Egln2* and *Axl* and retains subfamily order in *Neotoma* and *Peromyscus*; gene order is also similar to that of other vertebrates. (**B**) A second gene island composed of variable numbers of *cyp2B* genes arose in the New World rodents only, and underwent further expansion in *N. lepida*. (**C**) A tree of the inferred functional genes from each species was used to determine gene cluster identity/orthology, and is color coded similarly across all figure panels. Numbers at each node indicate bootstrap values.

The pattern of expansion is even more remarkable for *cyp2B*, where copy number evolution among *Neotoma* spp. was previously documented using combined quantitative PCR and cDNA cloning approaches (Malenke et al. 2012). First, we infer the evolution of a novel gene island containing only tandemly arrayed *cyp2B* genes in New World rats and mice (*Neotoma* + *Peromyscus*). *cyp2B* genes are typically found between the single-copy *cyp2G* and *cyp2S* genes in the 2ABFGST gene array and not elsewhere. This is the case for *Mus musculus* as well as other mammals such as humans and other primates (Hoffman and Hu 2007; Thomas 2007). Comparisons of this gene array in the past have shown dynamic rearrangement via duplication, deletion, and inversion within a genomic region bounded by the *Egln2* and *Axl* genes, but noted that translocations were limited to within these bounds (Hu et al. 2008). New World rodents show similar dynamic rearrangement of *cyp2B* genes within the ancestral location of this array, evidenced by the 2 extra functional copies and several pseudogenes in *P. maniculatus* compared to the woodrats, and the inverted orientation of the *cyp2B* gene closest to *cyp2S* in the woodrats (Figure 3A). Here, we report for the first time that New World rodents evolved a novel gene island outside the bounds of the ancestral array, comprised solely of variable numbers of *cyp2B* genes (Figure 3B). Based on the relationships among these copies, the novel island likely resulted from a duplication and translocation of the *cyp2B* gene closest to *cyp2S* to a location more than 1 MB away on the same chromosome, bounded by genes encoding zinc-finger proteins (zfp), *Tescl*, and *Lypd5*.

The novel *cyp2B* gene island consists of duplicates of the sequences called *cyp2B37* by Wilderman and colleagues (Wilderman et al. 2014) (Figure S11), while their *cyp2B35* and *cyp2B36* appear to be genes contained within the bounds of the ancestral gene island. The position of these *cyp2B37* sequences (Figure 3B) is not all that is novel about them, as they show unique enzymatic activity and substrate binding relative to *cyp2B35* and *cyp2B36* (Wilderman et al. 2014). It is possible that translocation to this new position facilitated the process of neofunctionalization of these genes. Additionally, copy number variation between *N. bryanti* and *N. lepida* accompanies this novel genomic location (Figure 3B), with 2 functional copies and 10 functional copies plus 6 pseudogenes in the respective species. These findings, in combination with the known functional uniqueness of cyp2B37 proteins, are consistent with the hypothesis that evolution within unstable gene islands is a source of adaptive novelty in the arms-races between herbivores and the toxic plants they feed on (Thomas 2007).

The *cyp3A* gene family in these rodents told a similar story to *cyp2B*, where conservation of genes involved in basal biosynthetic function remain generally conserved and genes involved in xenobiotic metabolism evolve in dynamic fashion. Again, our analyses using *Neotoma* and *P. leucopus* revealed both copy number evolution and the origination of unique gene islands of putative detoxification genes. All *cyp3A* genes are found on a single chromosome (Chr. 12 in *R.* norvegicus, Chr. 23 in *Peromyscus* and our *Neotoma* assemblies). In *R. norvegicus*, two *cyp3A* gene islands are approximately 7 Mb apart (Figure 4A). The first is small, containing the *cyp3A9*-like genes. This island showed general conservation among rodent species, with two functional copies in *R. norvegicus*, two in *P. leucopus*, three in *N. bryanti*, and one plus two pseudogenes in *N. lepida*. Moreover, these genes clustered together in the gene phylogeny, and this degree of conservation is likely reflective of the role cyp3A9 genes play in steroid hormone biosynthesis.

**Figure 4:**
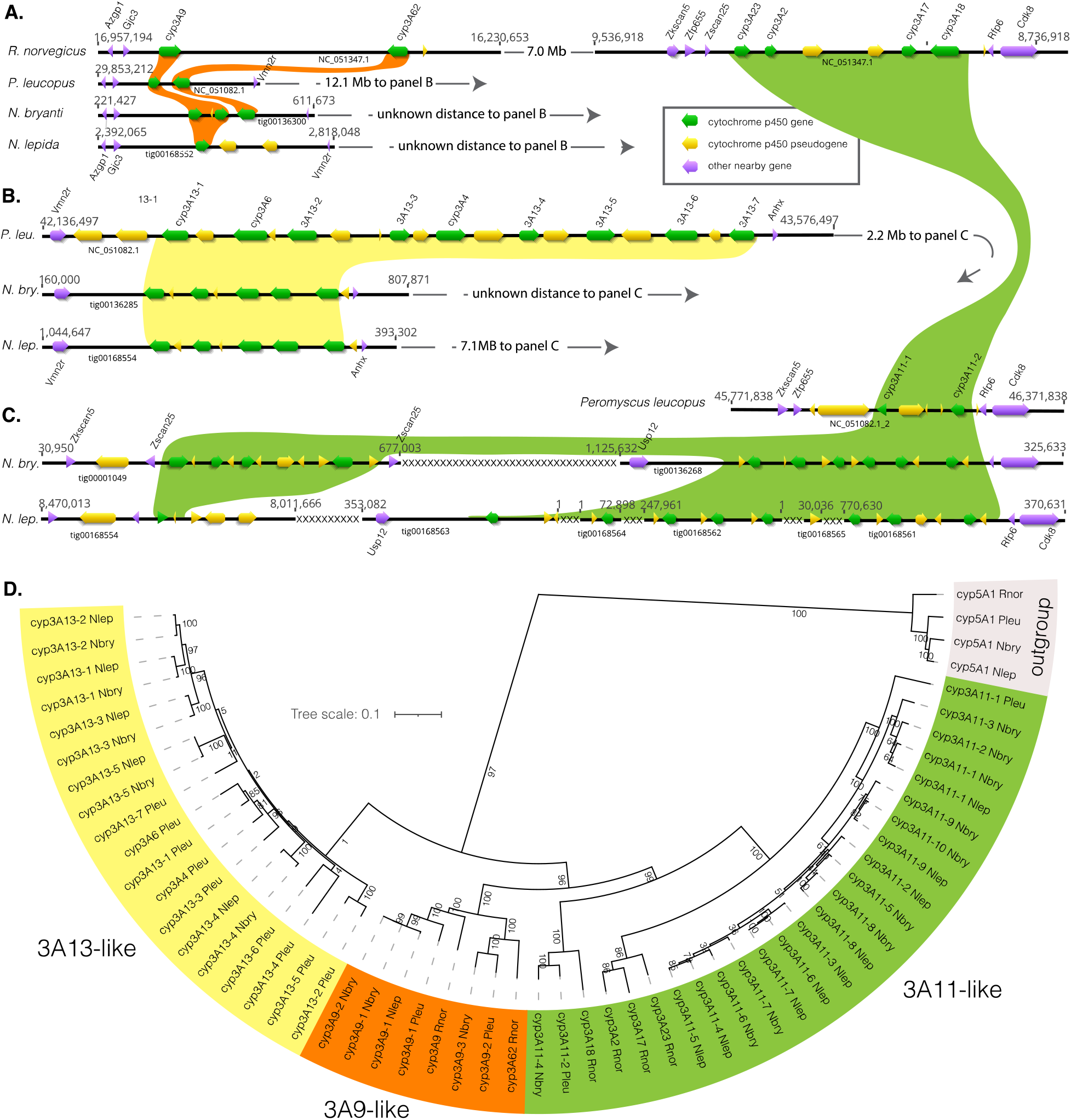
Genomic architecture and evolution of the *cyp3A* gene islands in four rodents. Visualization of the contigs containing these genes was based on existing annotations for the Norway rat (*Rattus norvegicus*; mRatBN7.2) and white-footed mouse (*Peromyscus leucopus*; UCI_PerLeu_2.1), and manual annotation of these regions for *Neotoma bryanti* and *N. lepida*. (**A**) Organization of *R. norvegicus cyp3a* genes on chromosome 12 (top contig), as well as the largely conserved *cyp3a9*-like genes found in *P. leucopus* and both *Neotoma*. (**B**) A second gene island has no clear orthology with a region in *R. norvegicus*. (**C**) The downstream *R. norvegicus* genes in (A) are related to a third downstream gene island in *P. leucopus* that is further divided into a third and fourth gene island in *Neotoma*. (**D**) A tree of the functional genes from each species was used to determine gene cluster identity/orthology, and is color coded similarly across all figure panels. Numbers at each node indicate bootstrap values.

In contrast, the remaining *cyp3A* gene islands all have evolved dynamically among the New World rodent species studied here. The second and final gene island in *R. norvegicus* contains four functional *cyp3A* genes and is bounded by several non-p450 genes that appear in the other taxa. However, between *cyp3A* islands syntenic with this second *R. norvegicus* island lies another island that appears unique to the New World rodents (Figure 4B). This array is bounded by *Vmn2r* and *Anhx*, and contains nine functional copies of *cyp3A* in *P. leucopus* (predominantly named *cyp3A13* in the RefSeq annotation) and 5 in each woodrat. We used the *cyp5A1* genes from each genome to form an outgroup to help us understand the relationships among these gene islands and among species, and found that the island containing the *cyp3A13*-like sequences is the sister group of the other *cyp3A* genes (Figure 4D). As such, we conclude that this family must have been present in the ancestor of all the rodents studied herein, and must therefore have been subsequently lost in the lineages leading to *R. norvegicus*.

On *Peromyscus* chromosome 23, we encountered *cyp3A11*-like genes that are similar in their count and sequence to their putative orthologs in *R. norvegicus* (Figure 4C). However, this pattern changes drastically when the two *Neotoma* species are added to the comparison: the inferred ancestral single gene island is instead two separate islands separated by the non-p450 gene *Usp12*. The novel fourth *cyp3A* gene island in *Neotoma* appears to have arisen, at least in part, due to partial duplication and inversion of a large segment of DNA. The *Zscan25* gene, which bound one side of this gene island in both *R. norvegicus* and *P. leucopus*, appears on each end of the novel island of *cyp3A11*-like genes in *N. bryanti* (Figure 4C, left side), and a large *cyp3A* pseudogene now separates *Zkscan5* and *Zscan25* on the 5’ side of the array. Altogether, these duplicated genes and entire new gene islands account for a large increase in *cyp3A* gene counts in the New World rodents compared to the 6 functional copies of *cyp3A* in *R. norvegicus* (Table 2), with the woodrats again having the largest number of total functional copies among the species examined.

## Conclusions

Our study contributes the first nearly complete, thoroughly annotated genomes of the genus *Neotoma*. These genomes provide the basis for understanding aspects of adaptive evolution in these and related species, and in particular, the dynamic evolution of the p450 gene family involved in detoxification and diet-related adaptations. These new genomic resources lay the groundwork for future studies into the adaptability of this unique genus (and vertebrate herbivores in general), and the success and efficiency of the trio binning method should encourage other researchers studying hybridizing species to consider utilizing this approach.

## Data availability

The CLC assemblies, Canu assemblies, chromosome scaffolds and accompanying annotations for *Neotoma bryanti* and *Neotoma lepida* have been deposited at the Open Science Framework and are available at https://osf.io/xck3n/.

## Acknowledgments

We thank Dr. Richard Clark for generously sharing his computational resources with us for this work and Sebastian Smith for assistance with UNR’s Pronghorn. We are also indebted to Dr. James Patton for providing the male *Neotoma lepida* used for the F_1_ cross, Madeline Nelson for performing the animal husbandry, Jennifer Dixon and Kika Kitanovic for carrying out RNA preparation for the transcriptome work, Whitney Kenner for preparing the DNA samples, and Danny Nielson for initial genotyping to confirm the identity of genome animals. We are grateful to Dylan Klure for his comments on the manuscript and Dr. Sara Weinstein (juniper and creosote images), Maggie Doolin (*N. bryanti* image) and Dr. Kevin Kohl (*N. lepida* image) for the photographs in Figure 1. This research was supported by grants from the National Science Foundation to M.D.M. (IOS-1457209 and OIA-1826801) as well as M.D.D. and M.D.S. (IOS-1656497).

