## Supplemental Figures S1-S11 for "Trio-binned genomes of the woodrats *Neotoma bryanti* and *N. lepida* reveal novel gene islands and rapid copy number evolution of xenobiotic metabolizing cytochrome p450 genes"

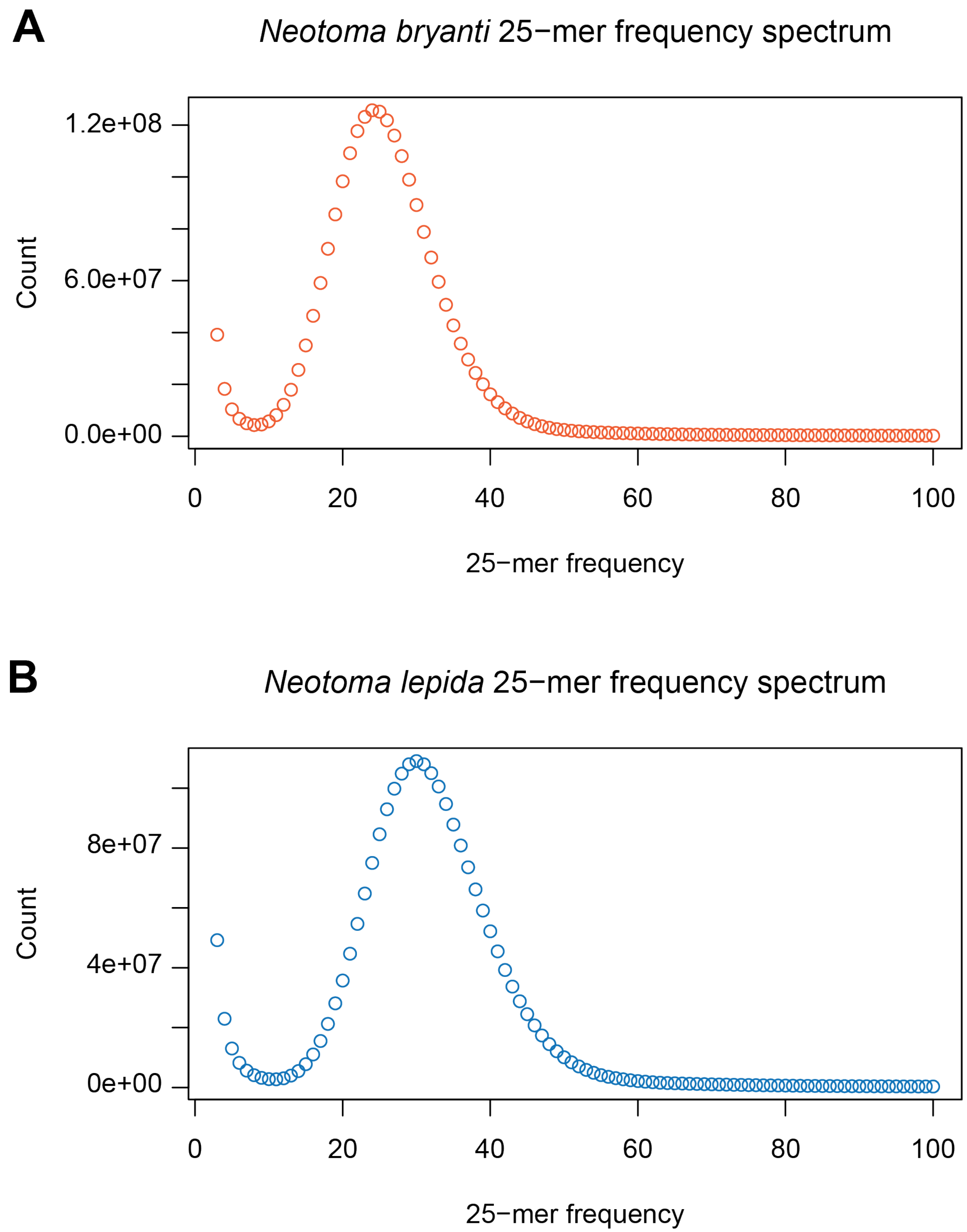


**Figure S1: 25-mer frequency spectra for the partitioned *Neotoma bryanti* and *Neotoma lepida* reads.** 25-mer spectra counts generated by Jellyfish 2.3.0 are shown for **(A)** *N. bryanti* and **(B)** *N. lepida*.

**Figure S2:** ***Neotoma bryanti*** **contigs aligned against** ***Peromyscus leucopus*** **chromosome scaffolds.**

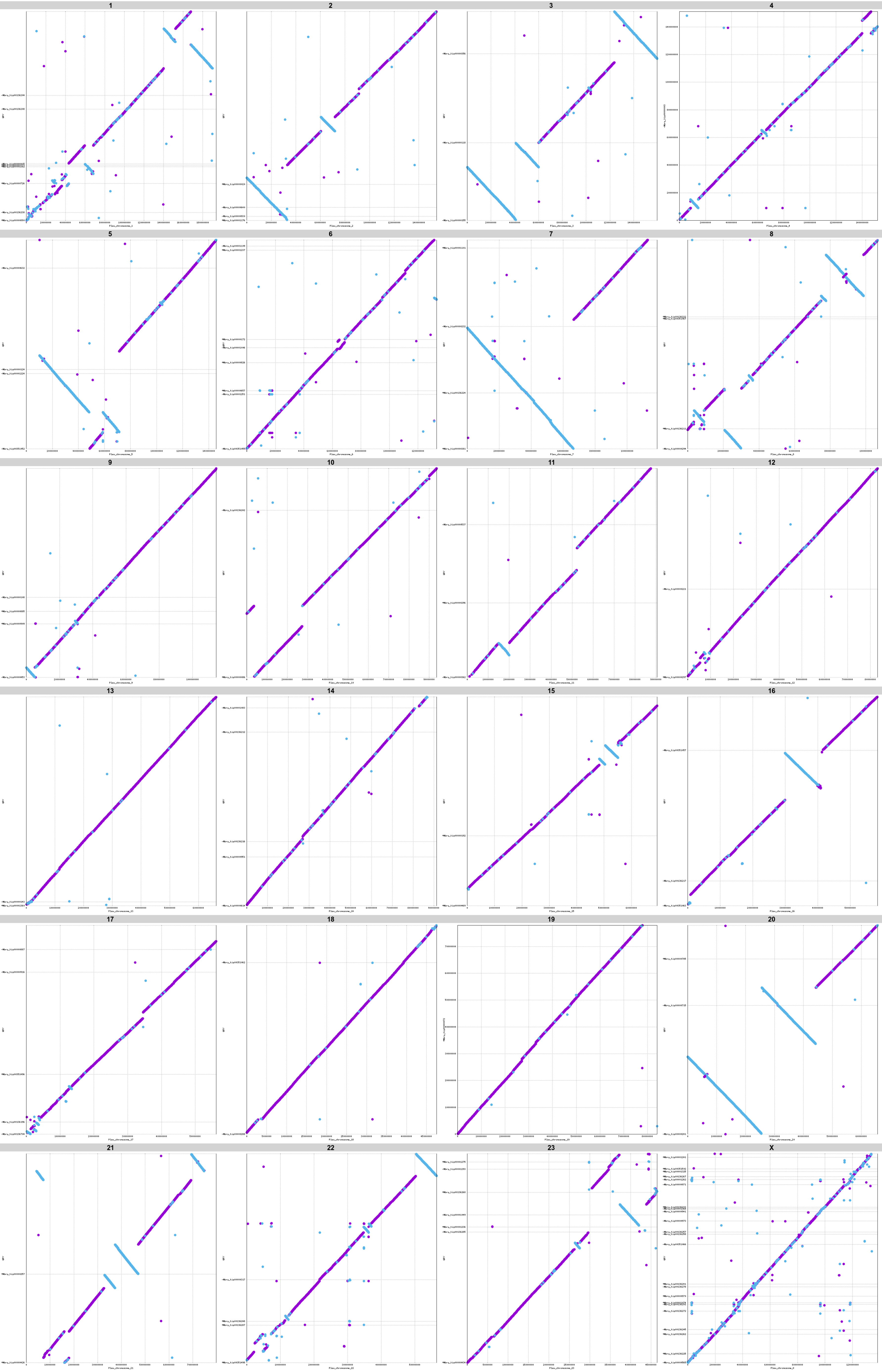

**Figure S3:** ***Neotoma bryanti*** **contigs aligned against** ***Peromyscus maniculatus*** **chromosome scaffolds.**

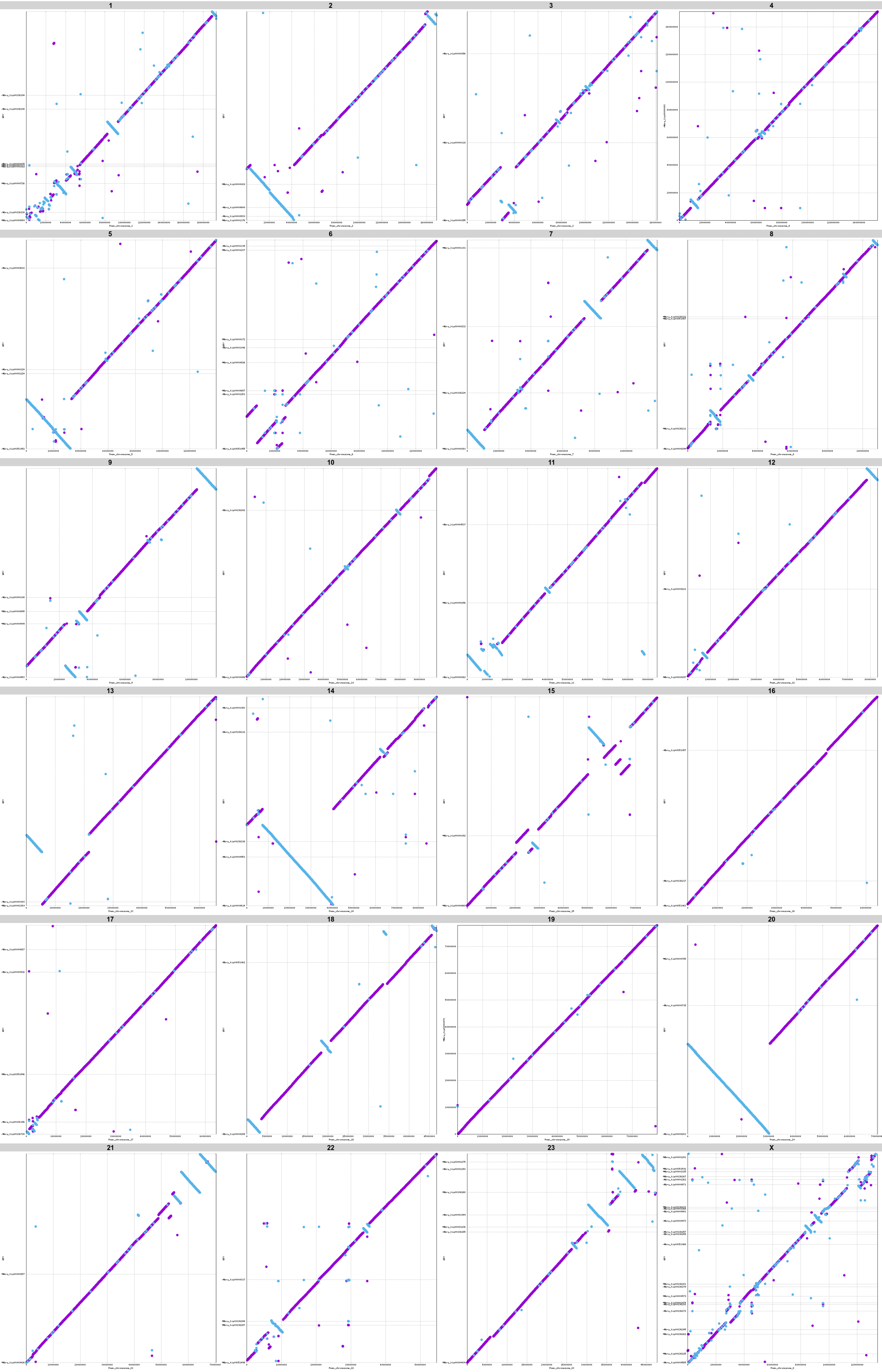

**Figure S4: *Neotoma bryanti* contigs aligned against *Peromyscus nasutus* chromosome scaffolds.**

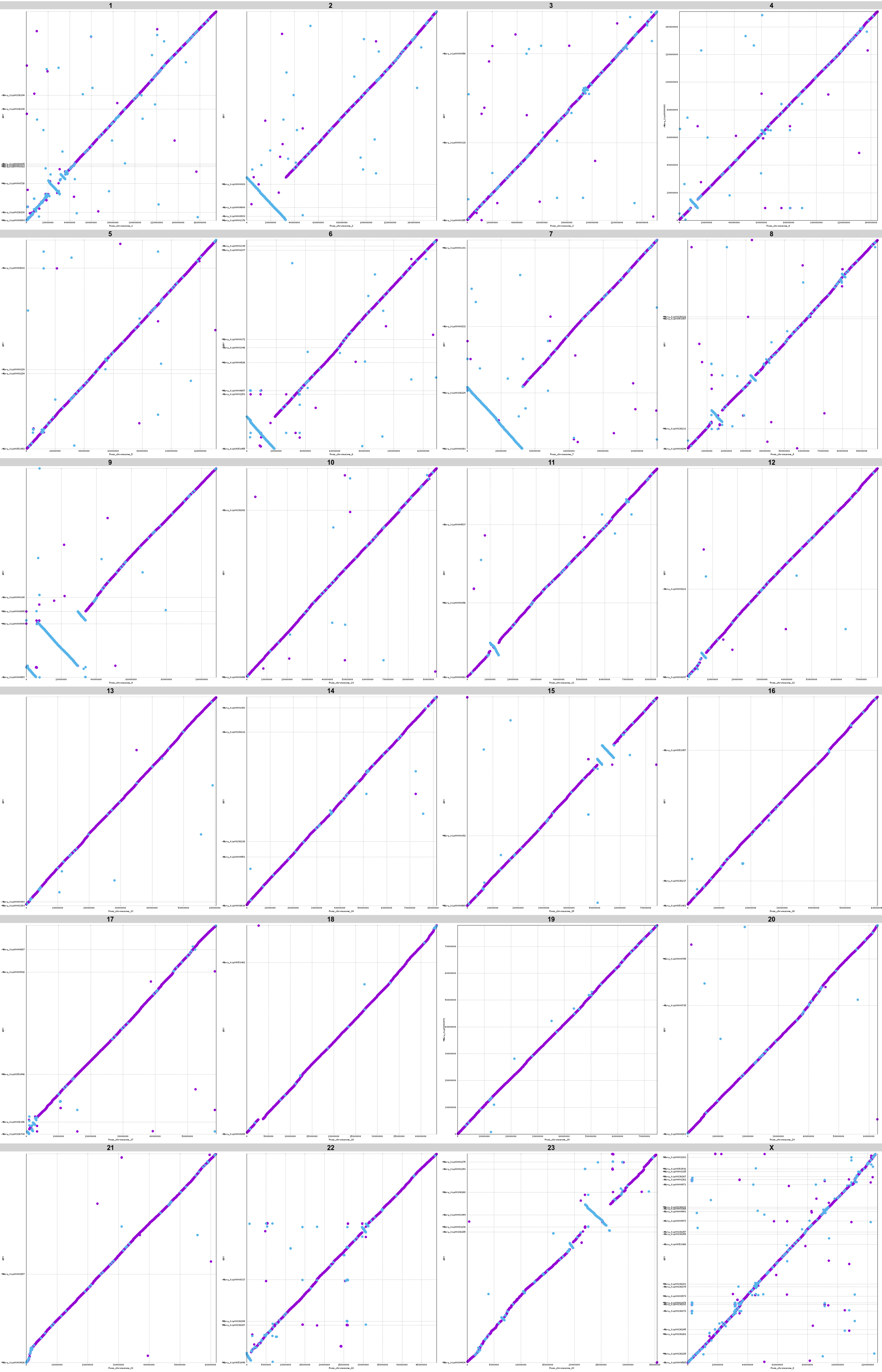

**Figure S5:** ***Neotoma lepida*** **contigs aligned against** ***Peromyscus leucopus*** **chromosome scaffolds.**

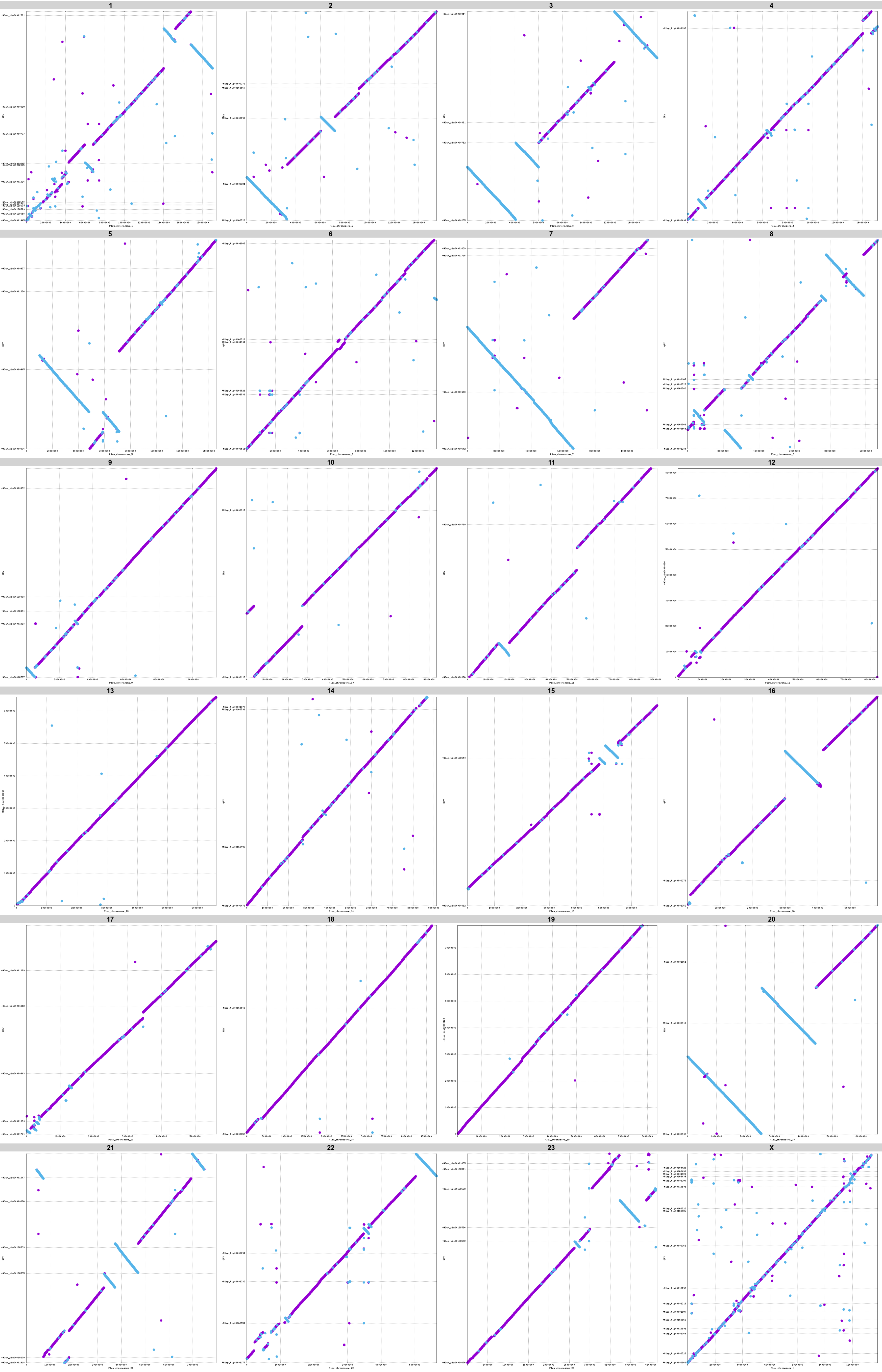

**Figure S6:** ***Neotoma lepida*** **contigs aligned against** ***Peromyscus maniculatus*** **chromosome scaffolds.**

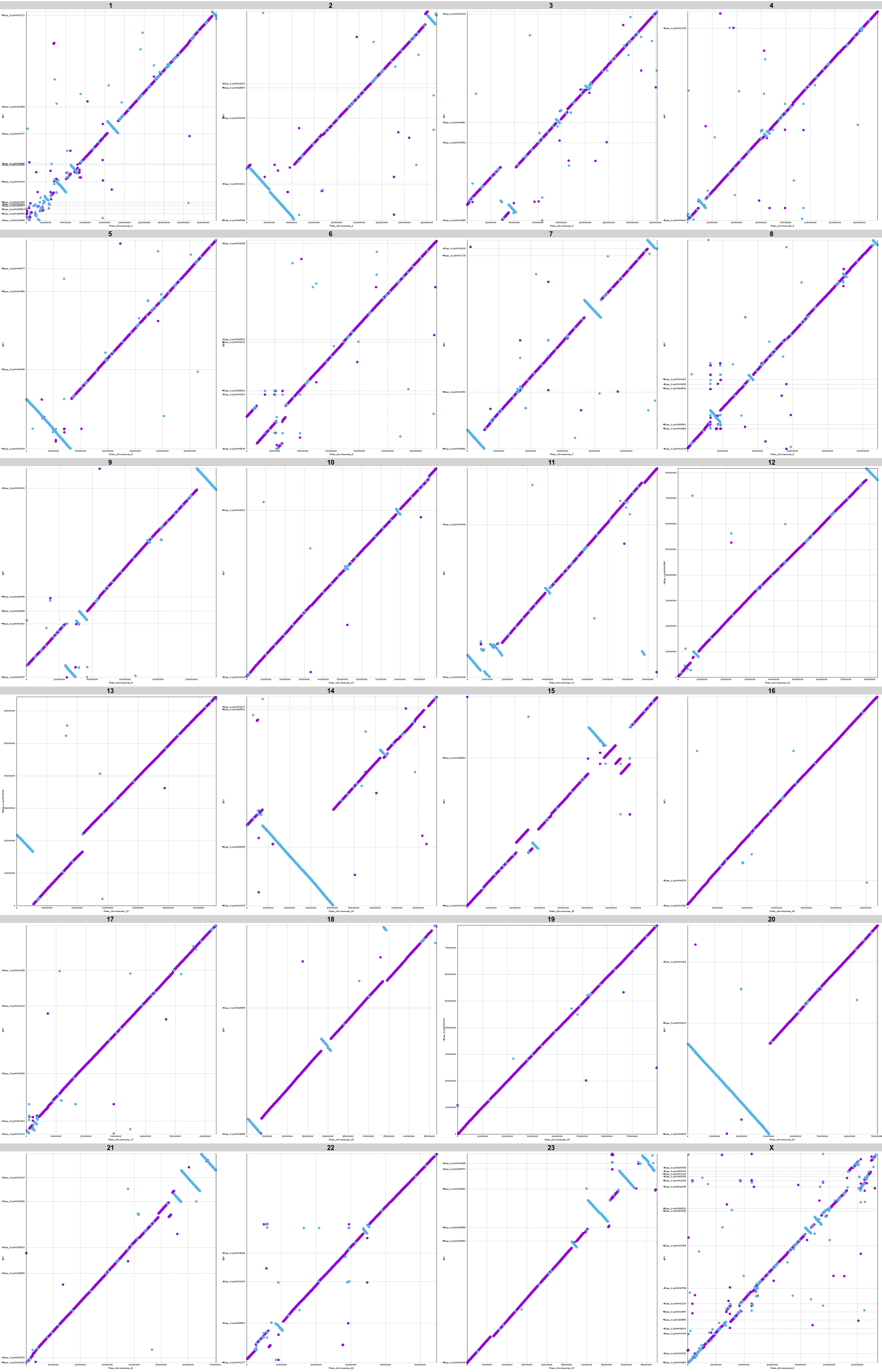

**Figure S7:** ***Neotoma lepida*** **contigs aligned against** ***Peromyscus nasutus*** **chromosome scaffolds.**

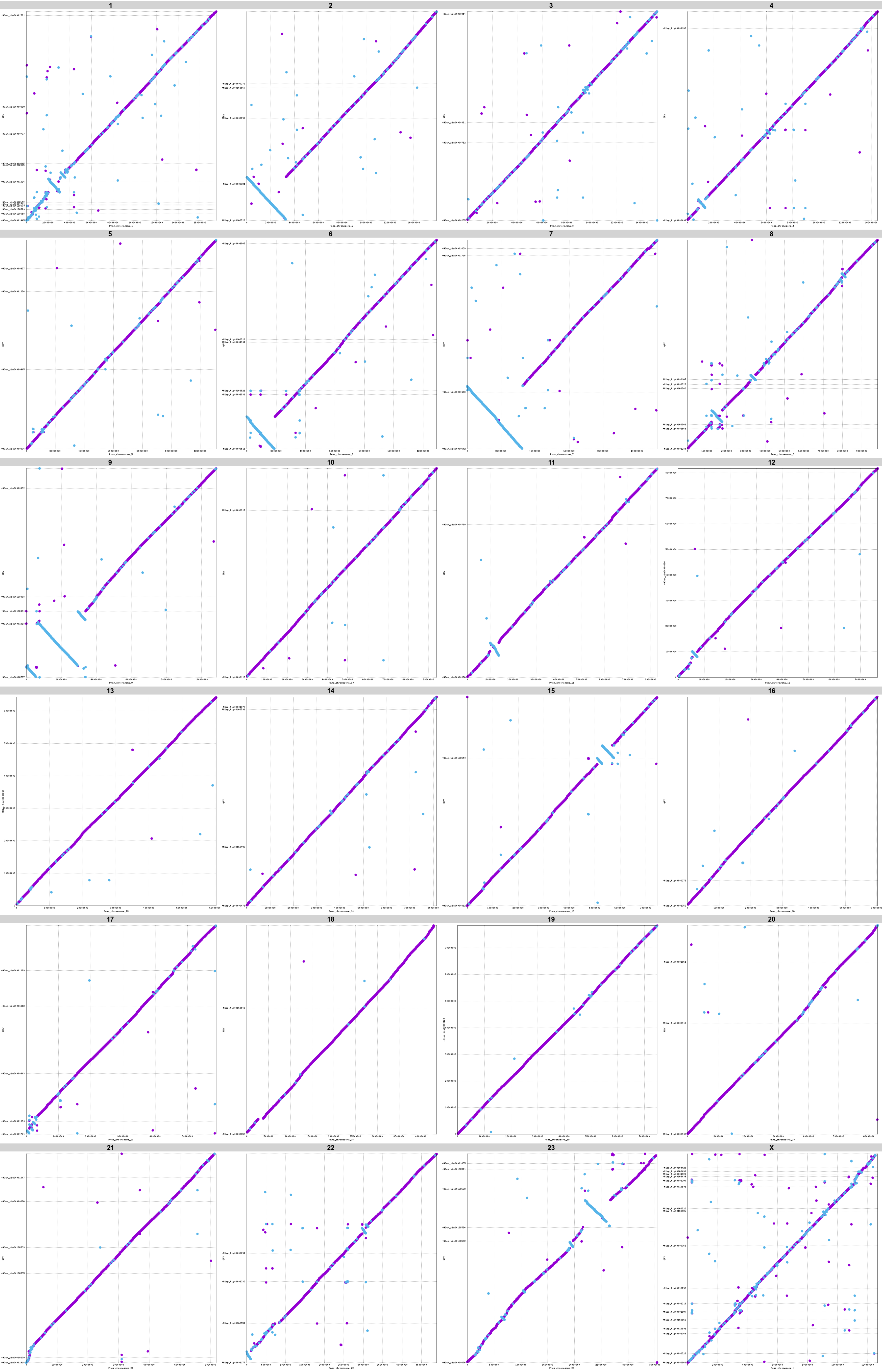

**Figure S8:** ***Neotoma bryanti*** **contigs aligned against** ***Neotoma lepida*** **chromosome scaffolds.**

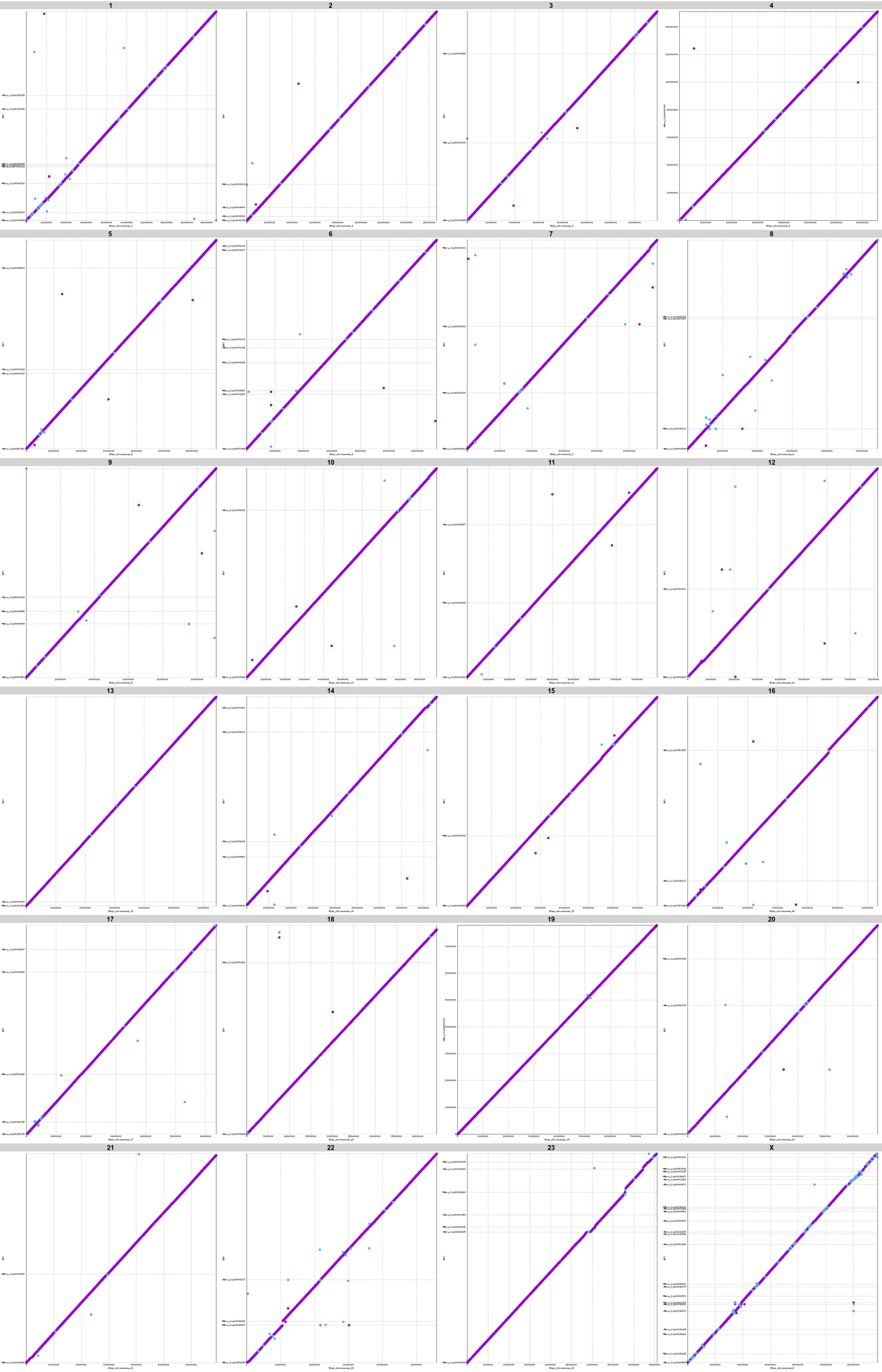

**Figure S9: *Neotoma lepida* contigs aligned against *Neotoma bryanti* chromosome scaffolds.**

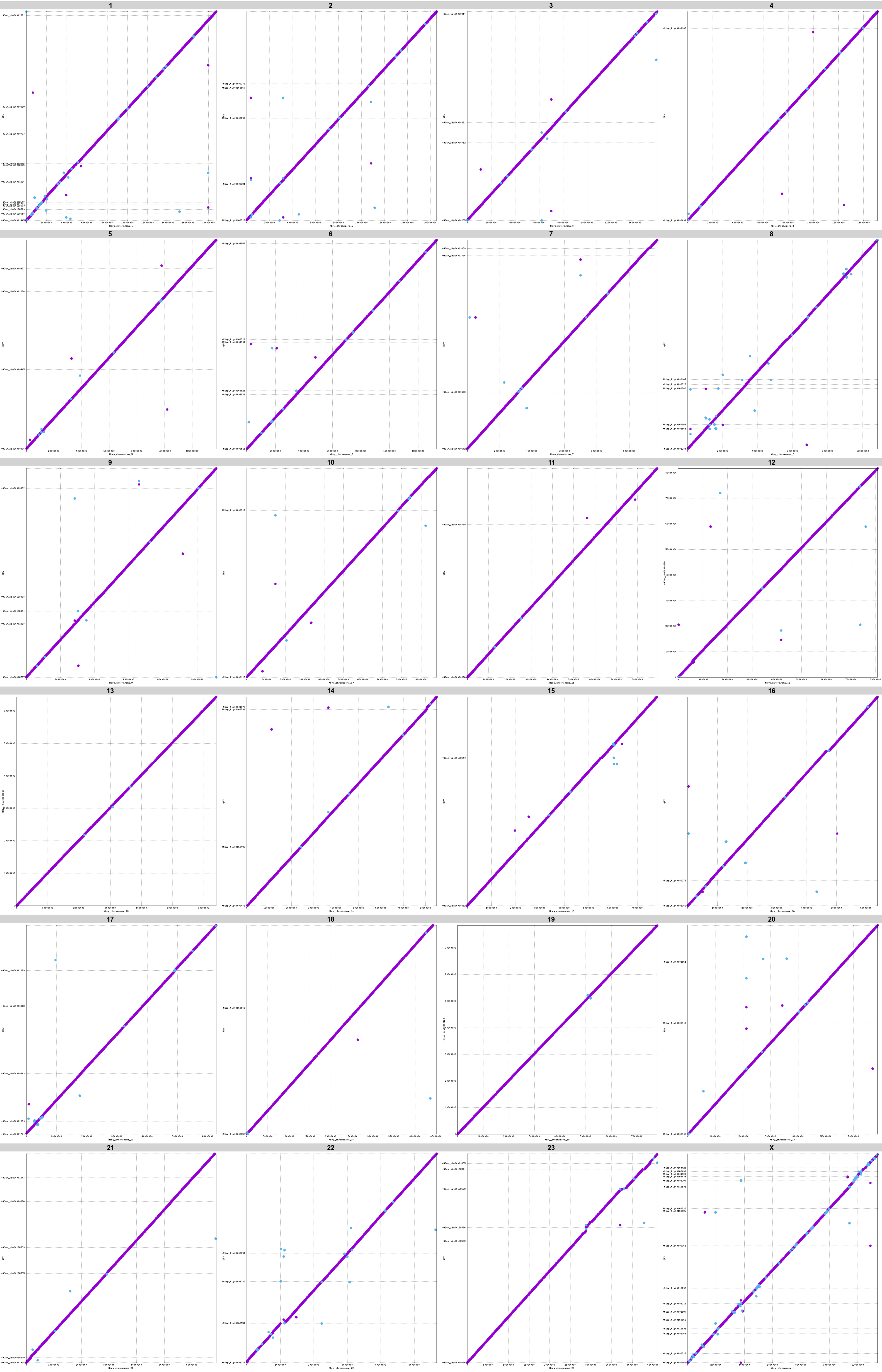

**Figure S10: Alignments of assemblies used to resolve the CYP2 region in *N. lepida*.** Alignments of the (**A**) 6.58 Mb and 11.98 Mb contigs from the 40× assembly and (**B**) 13.63 Mb contig from the all reads (AR) assembly against the 18.55 Mb CYP2 sequence produced by merging the three contigs with quickmerge 0.3. Note that the 13.63 Mb contig is composed of sequence contained in both contigs from the 40× assembly. (**C**) Alignment of the four CYP2 contigs from the *N. lepida* Canu assembly that appear to compose the entirety of this sequence. (**D**) Alignment of the remaining three CYP2 contigs from the Canu assembly that appear to be potential repetitive sequences or assembly errors. These contigs align to the same region as that occupied by Nlep_tig00418827 (top).

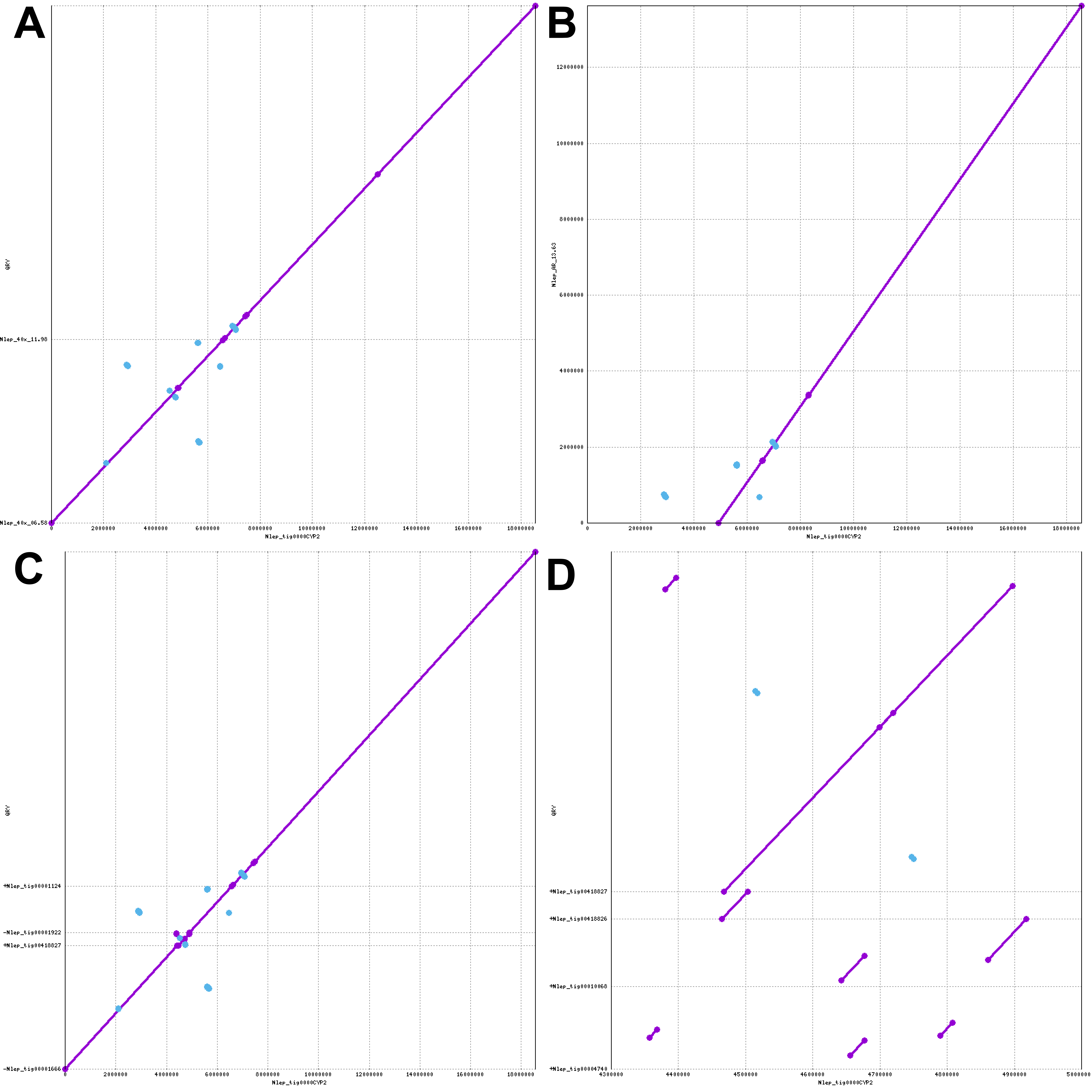

**Figure S11: Phylogeny of cyp2ABFGST genes.** Unrooted phylogeny of the cyp2ABFGST genes from the four rodent species studied, as well as protein sequences of the genes named cyp2B35, cyp2B36, and cyp2B37 from Wilderman et al. (2014).

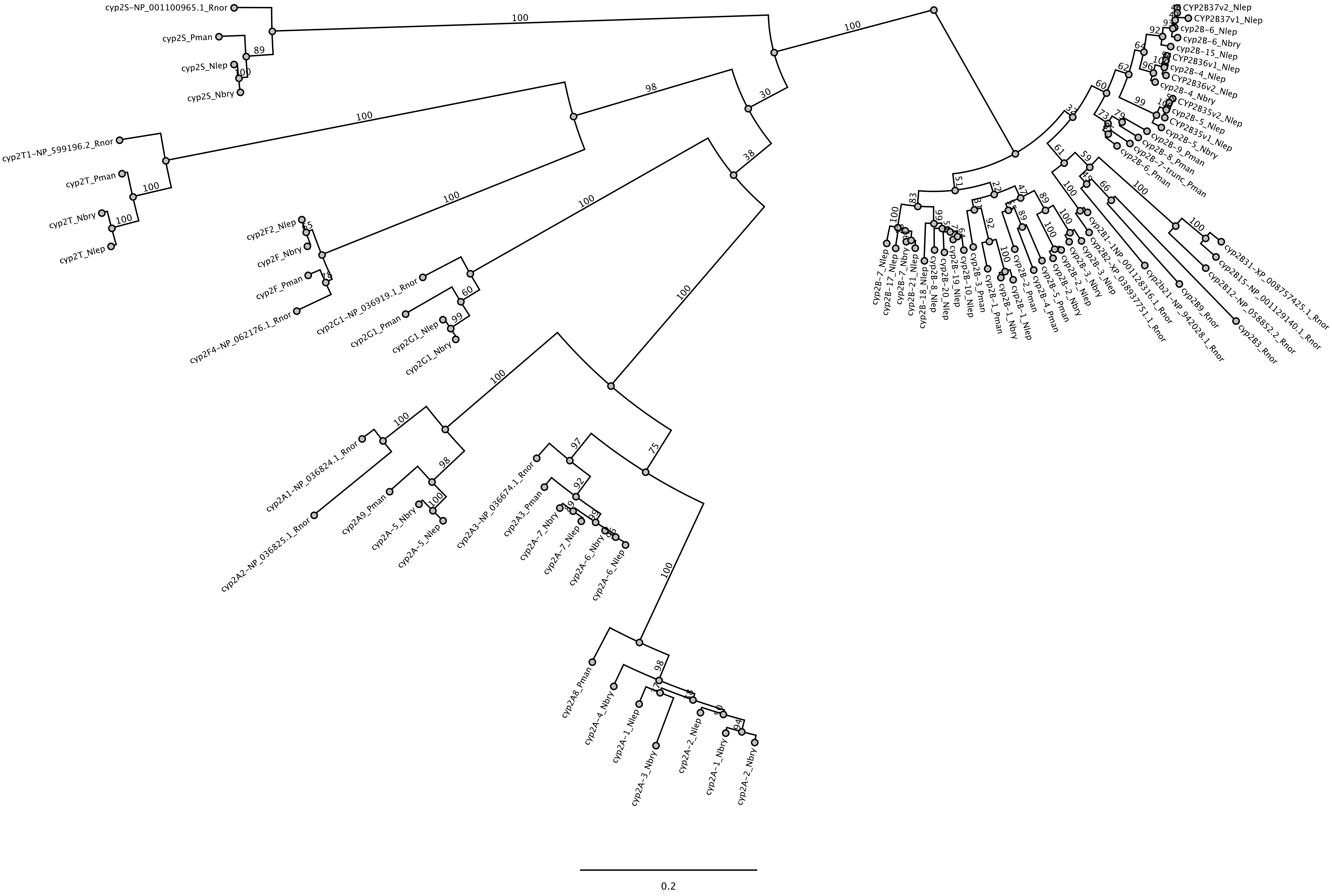
